# A central helical fulcrum in eIF2B coordinates allosteric regulation of Integrated Stress Response signaling

**DOI:** 10.1101/2022.12.22.521453

**Authors:** Rosalie E Lawrence, Sophie Shoemaker, Aniliese Deal, Smriti Sangwan, Aditya Anand, Lan Wang, Susan Marqusee, Peter Walter

## Abstract

The Integrated Stress Response (ISR) enables cells to survive a variety of acute stresses, but chronic activation of the ISR underlies age-related diseases. ISR signaling down-regulates translation and activates expression of stress-responsive factors that promote return to homeostasis, and is initiated by inhibition of the decameric guanine nucleotide exchange factor eIF2B. Conformational and assembly transitions regulate eIF2B activity, but the allosteric mechanisms controlling these dynamic transitions are unknown. Using hydrogen deuterium exchange-mass spectrometry and cryo-EM, we identified a single alpha-helix whose orientation allosterically controls eIF2B conformation and assembly. Biochemical and signaling assays show that this “Switch-Helix” controls eIF2B activity and signaling in cells. In sum, the Switch-Helix acts as a fulcrum of eIF2B conformational regulation and is a highly conserved actuator of ISR signal transduction. This work uncovers a novel allosteric mechanism and unlocks new therapeutic possibilities for ISR-linked diseases.

## Introduction

The integrated stress response (ISR) is a conserved signaling network that promotes cellular fitness in response to biological stress by reprogramming translation and metabolism^1,2^. Acute activation of the ISR down-tunes general protein synthesis, while selectively activating expression of stress-responsive factors including chaperones, redox balancers, and amino acid importers^2^. When a return to homeostasis is not achieved, ISR signaling can activate apoptosis^3^. While acute activation of the ISR is adaptive, a chronic ISR signaling state is associated with a wide range of age-related and neurodegenerative pathologies, including Alzheimer’s disease, brain injury-induced dementia, cancer, and Down syndrome^4–10^.

The ISR is carried out by a complex circuitry that defines downstream signaling by controlling the rate of translation initiation complex assembly^11^. The rate-determining step in this pathway is carried out by eIF2B, a sophisticated heterodecameric guanine nucleotide exchange enzyme whose activity is controlled by its conformational state as well its assembly from sub-complexes (**Fig. 1a**)^12–14^. An important goal is to uncover the molecular mechanisms that enable these dynamic transitions. For instance, how does binding of modulators communicate allosterically across a large multisubunit complex to regulate signaling? Defining molecular mechanisms of eIF2B control will illuminate both basic principles of allosteric regulation of large signaling complexes as well specific network features of this core signaling pathway, and will open new avenues for therapeutic development.

**Fig. 1.**
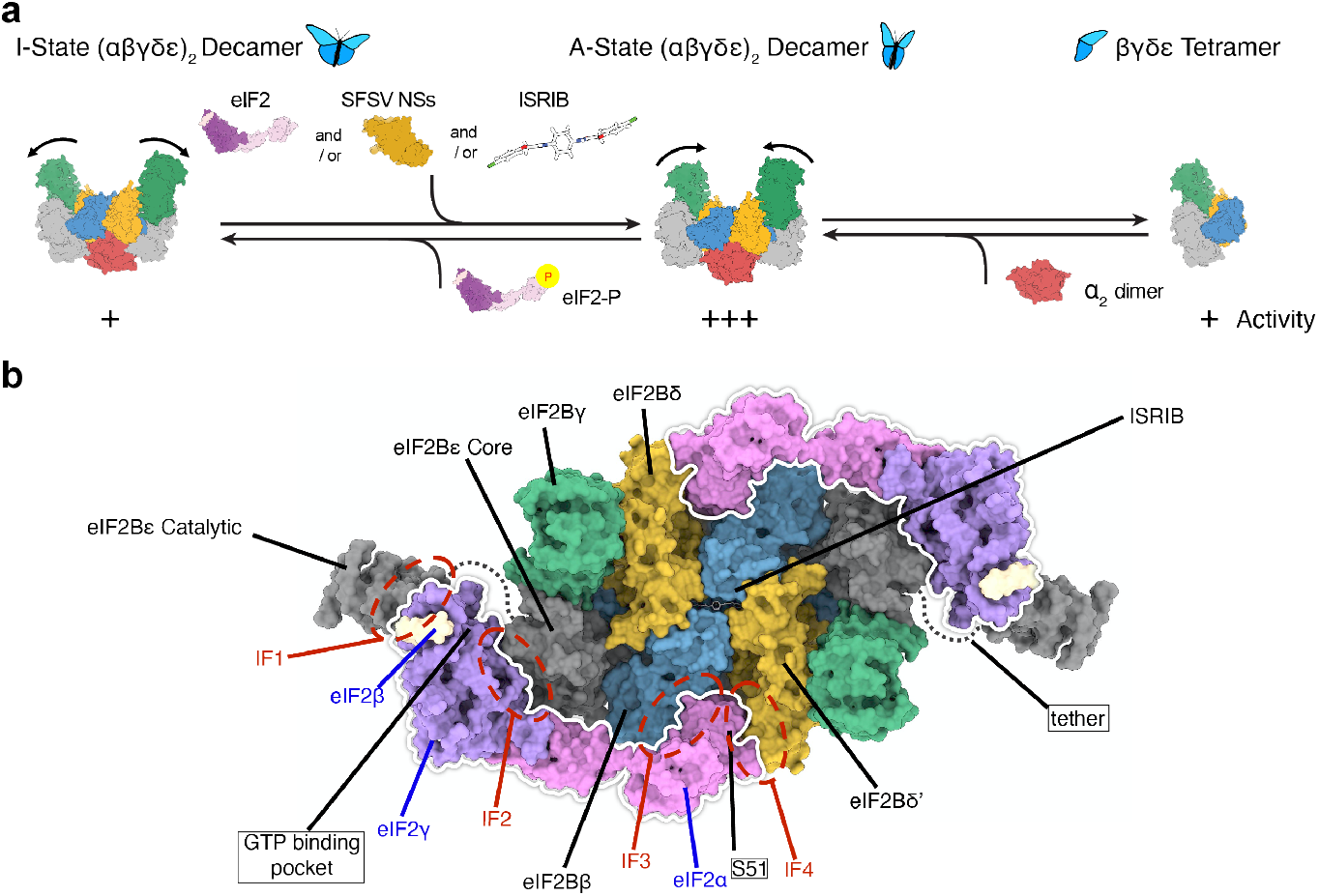
eIF2B activity is controlled by conformation and assembly state. (**a**) eIF2B is regulated by assembly state and conformation. Assembly from less-active tetramers (right) into more-active decamers (middle) is driven by availability of the eIF2Bα_2_ dimer. The less-active I-state conformation (left) is driven by eIF2-P binding. The active A-state conformation (middle) is driven by eIF2, NSs, or ISRIB binding. (**b**) A surface representation of a model of two eIF2 trimers and ISRIB bound to an eIF2B(αβγδε)_2_ decamer is shown. Individual subunits of eIF2 and eIF2B are indicated. The eIF2 trimers are outlined in white and the locations of interfaces IF1-4 are indicated, as are the positions of eIF2 S51, the GTP binding pocket (empty in the structure), and ISRIB (shown in stick presentation). The eIF2Bα_2_ dimer is hidden in this orientation. eIF2Bε contains two domains including the eIF2Bε catalytic HEAT domain that is linked by a flexible tether which was not resolved in the structure.

eIF2B promotes GTP-loading of the trimeric GTPase eIF2 (composed of α, β, and γ subunits)^15^. This rate-limiting step enables eIF2 to assemble with methionyl tRNA to form the translation initiation factor ternary complex^16^. When ternary complex levels are low, translation of most mRNAs is inhibited, while a 5’ untranslated region (5’ UTR)-regulated mechanism promotes expression of stress-adaptive programs via selective translation of a few mRNAs including that encoding the transcription factor ATF4^11,15,17^. We define the ISR as the events triggered in cells that contain limiting ternary complex.

eIF2B is a central hub of the complex ISR signaling pathway^18,19^. Cellular stress is sensed by four distinct kinases–PERK, HRI, GCN2, and PKR–which each phosphorylate Serine 51 of eIF2α in response to misfolded proteins, redox and mitochondrial stress, amino acid deficiency, and viral infection, respectively^1^. Phosphorylation activates the ISR by converting eIF2 from a substrate of eIF2B (‘eIF2 substrate’) into its inhibitor (‘eIF2-P inhibitor’), impeding eIF2 GTP-loading and hence ternary complex assembly^20^. Hence, eIF2B is a highly tunable regulator of ISR outcome, and eIF2B activity is inversely proportional to ATF4 induction^18,19^.

The eIF2B complex is a two-fold symmetric heterodecamer composed of two copies each of subunits α, β, γ, δ, and ε^21–23^. eIF2B activity is controlled by two distinct structural changes: first, eIF2B activity is regulated by assembly state. The eIF2B complex can disassemble into two tetramers composed of subunits β, γ, δ, and ε (herein referred to as ‘eIF2B tetramer’ or ‘eIF2Bβγδε’) and one dimer composed of two α subunits (herein referred to as ‘eIF2Bα_2_’ or ‘α_2_ dimer’) (***Fig 1a***)^18,19,21^. eIF2B tetramers have reduced activity relative to fully assembled eIF2B decamers (later referred to as ‘eIF2B(αβγδε)_2_’ or ‘eIF2B decamers’). Thus, cells can tune ISR activity by regulating eIF2Bα_2_ availability, controlling the amount of fully active eIF2B decamers^14^. The small molecule ISRIB inhibits the ISR by binding across the tetramer-tetramer interface to promote eIF2B decameric assembly and increase its activity^21,22,24,25^.

eIF2B activity is also controlled by conversion between active and inactive conformations^13,14^. The inhibitor eIF2-P prompts conformational change in eIF2B by binding either of two (symmetrically identical) pockets between the eIF2Bα and eIF2Bδ subunits. This induces a complex-wide rocking motion in which the two symmetric eIF2B halves rotate away from each other, akin to the flapping of a butterfly’s wings (***Fig. 1a***). In turn, this widens the pockets where eIF2 substrates bind, located between eIF2Bβ and eIF2Bδ’, reducing substrate engagement and eIF2B activity (see ***Fig. 1b***). We refer to the inhibited “wings down” state as the “I-State”, and the active “wings up” state as the “A-State” (***Fig. 1a***)^14^. In the absence of binding partners, cryo-EM data suggests that eIF2B primarily samples the A-state^14^. The eIF2B A-State can be further stabilized by binding to eIF2 substrate, ISRIB and its analogs, or the viral ISR-inhibiting protein NSs^26,27^. The two states are negatively coupled: the A-State and I-State have alternatively accessible binding sites, such that the A-State has a properly formed eIF2-binding pocket, and the I-State has a properly formed eIF2-P-binding pocket^13,14^. Furthermore, the I-State eIF2B decamer has an activity that approximates that of eIF2B tetramers, which we previously posited is because the I-State rocking motion removes one of four eIF2 interaction interfaces, (labeled Interface 4 in ***Fig. 1b***)^14^.

We do not understand the molecular mechanism of eIF2B conformational remodeling. Specifically, it is unknown how binding of activators or inhibitors at distant sites coordinates allosteric remodeling of the eIF2B complex’s conformation across multiple subunits, ultimately controlling eIF2B activity and ISR signaling. Thus, it has remained an important goal to uncover the mechanisms that enable these dynamic transitions. Here, we use hydrogen-deuterium exchange monitored by mass spectrometry, cryo-EM, and biochemistry to map the allosteric mechanism controlling eIF2B activity. We propose a comprehensive allosteric mechanism that regulates both eIF2B conformational and assembly transitions, centered around our discovery of the eIF2B Switch-Helix. Our work unlocks new strategies to study and design therapeutics for the broad range of diseases characterized by chronic ISR signaling.

## Results

### Hydrogen Deuterium Exchange probes eIF2B conformational and assembly dynamics

To define molecular mechanisms of eIF2B’s dynamic regulatory transitions, we used hydrogen deuterium exchange-mass spectrometry (HDX-MS) to obtain a comprehensive profile of eIF2B’s structural flexibility. HDX-MS monitors the exchange of protein backbone amide hydrogens for deuterium atoms from a deuterated buffer at the resolution of small peptides^28^. An individual amide’s rate of exchange is directly related to its solvent accessibility and local stability, and thus reports on changes in structure and conformational freedom^29–31^. HDX-MS experiments are carried out by exposing protein to deuterated solvent (***Fig. 2a, Step 1***), quenching the deuteration reaction at various timepoints (via low pH and low temperature), proteolyzing the sample under quenched conditions (***Step 2***), followed by in-line liquid chromatography-mass spectrometry (***Step 3***) to detect the number of deuterons per peptide at each timepoint. The changes in mass due to uptake of deuterons can then be mapped onto the structure, revealing time-resolved structure and conformational freedom (***Step 4***).

**Fig. 2.**
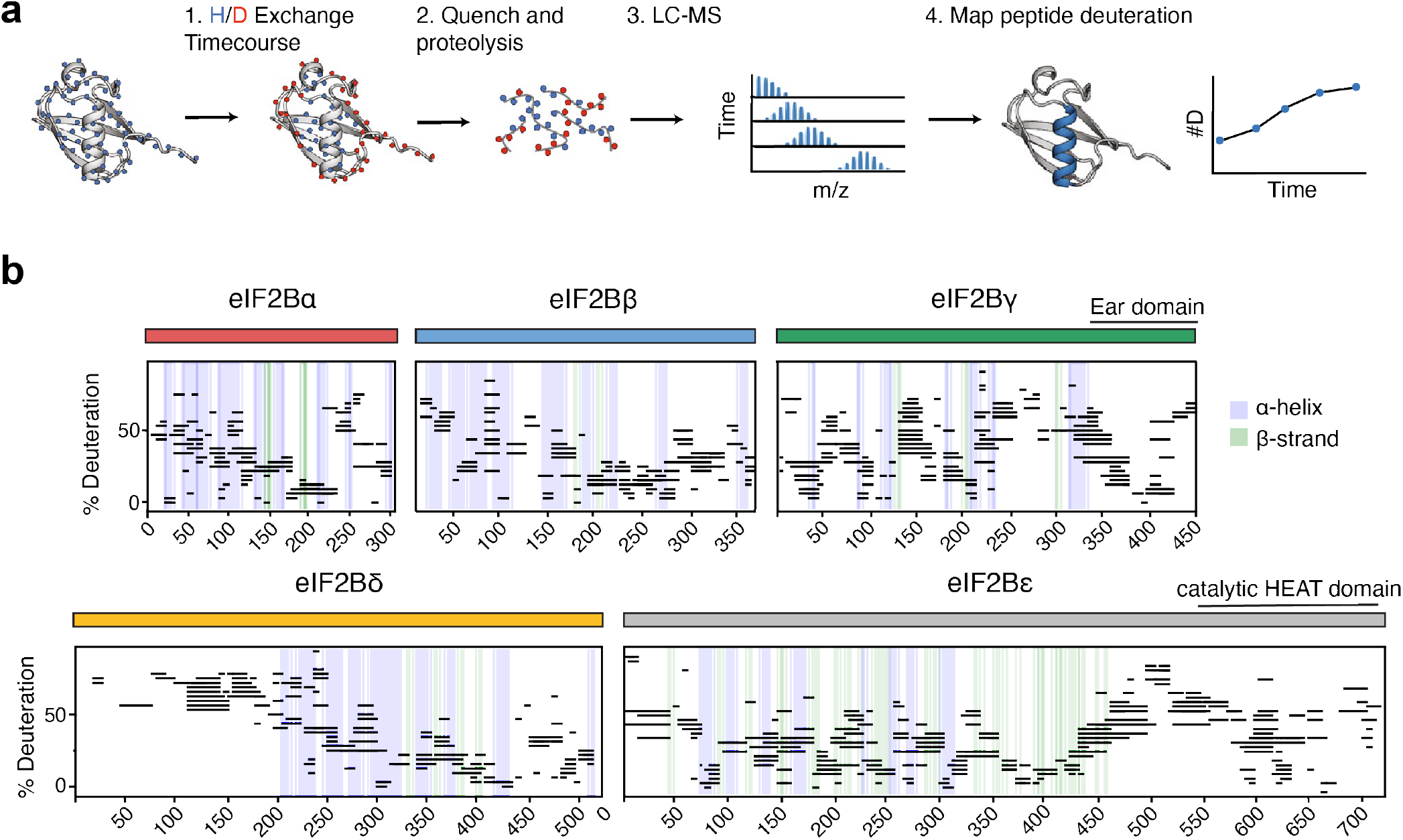
HDX-MS probes eIF2B structure. (**a**) Schematic of an HDX-MS experiment. 1. Protein is incubated in deuterated solvent and solvent-exposed amino acids are allowed to exchange with deuterium until defined timepoints. 2. Exchange is quenched via pH drop and protein is protease digested. 3. Peptide deuteration is detected via liquid chromatography-mass spectrometry (LC-MS). 4. Peptide deuteration uptake is plotted over time, and interpretated in the context of structural information. (**b**) Percent deuteration after 100 seconds of deuterium labeling for every peptide in one apo eIF2B dataset. Solid colored bars indicate each eIF2B subunit, corresponding to color scheme in Fig. 1. Each horizontal line represents an individual peptide spanning the residues indicated on the x axis, with percent deuteration (neglecting back exchange) indicated on the y axis. α-helices are indicated in blue vertical lines, and β-strands are indicated in green vertical lines, derived from apo eIF2B structure PDB 7L70. The eIF2Bγ C-terminal “Ear domain” and the eIF2Bε C-terminal Heat domain (regions not well characterized by Cryo-EM) are indicated.

After extensive optimization of protease digestion and quench conditions, we obtained excellent sequence coverage (92%) of the large eIF2B decamer (2369 unique amino acids). This enabled monitoring of nearly its entire sequence space, including conformationally flexible regions not modeled in existing cryo-EM structures (all peptides we monitored showed EX2 behavior) (***Fig. 2b, Supp. Figs. 1–2***)^13,18,21,22,26,27^. We observed broad agreement between regions of HDX-MS protection and experimental secondary structure assignment based on the cryo-EM structure of apo-eIF2B (***Fig. 2b***, see blue and green vertical bars indicating α helices and β sheets, respectively)^14^.

### HDX-MS analysis of eIF2B A-State and I-State transitions identifies a remodeled helix

To uncover underlying mechanisms that control the eIF2B A→I-State conformational change and/or its eIF2Bβγδε→eIF2B(αβγδε)_2_ (i.e., tetramer→decamer) assembly, we performed comparative HDX-MS experiments. To this end, we biased the population of eIF2B decamers toward the A-State with the addition of the viral ISR-inhibitor NSs or the small molecule ISRIB-analog 2BAct and carried out HDX-MS to compare the time-dependent deuteration uptake (uptake profile) between the apo-state and the A-state (***Fig. 3a, Supp. Fig 3***, and ***Supp. Fig. 5a***). Similarly, we biased eIF2B decamers toward the I-State by the addition of eIF2-P, and again compared the uptake profile between the I-State and the apo-State (***Fig. 3b***). We defined a region as showing increased protection if it showed less deuterium uptake in the defined A- or I-States relative to the apo-State.

**Fig. 3.**
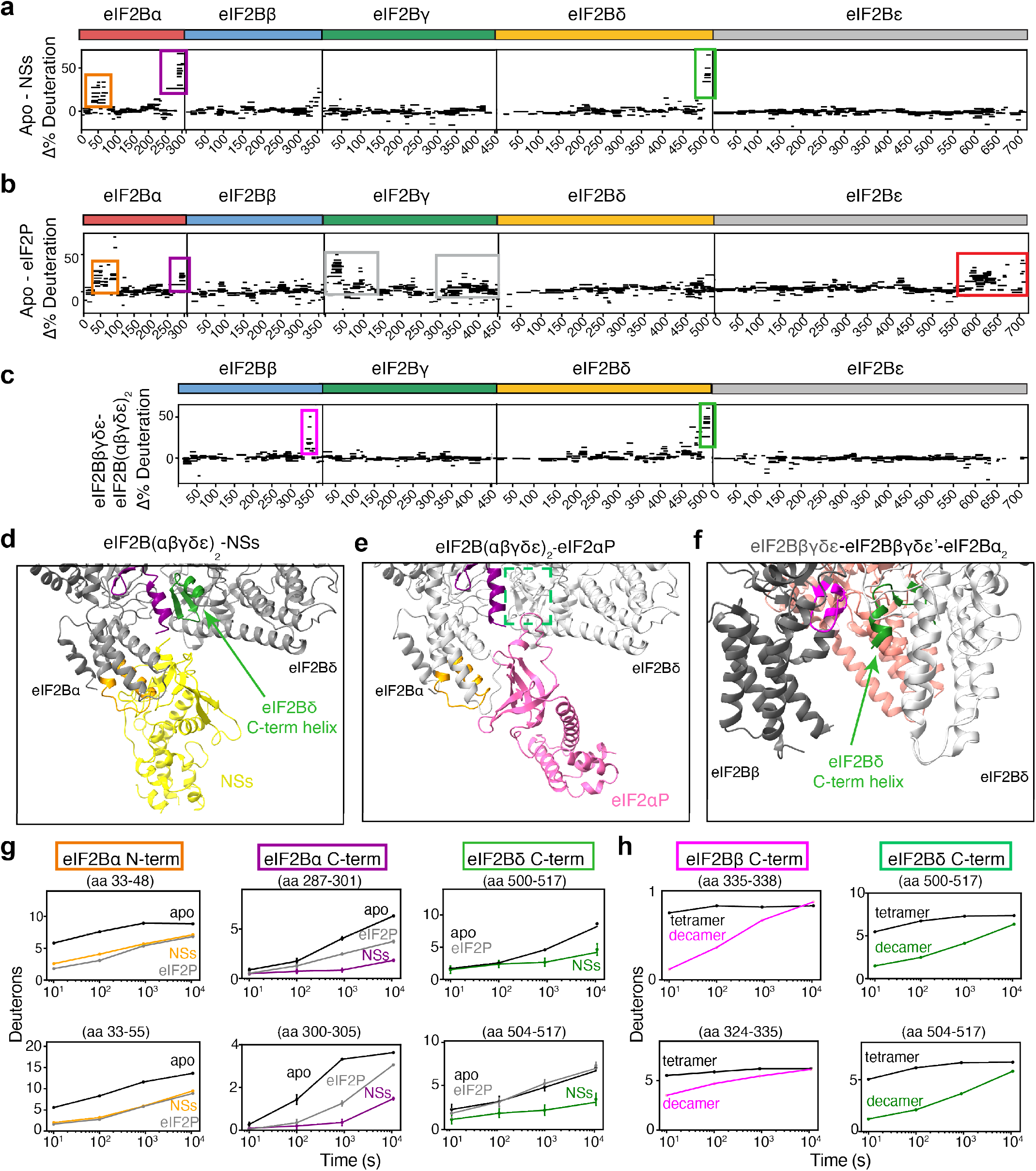
HDX-MS analysis of eIF2B conformation and assembly states identifies remodeling of the same helix. (**a-c**) Representative percent deuteration difference maps for (**a**) apo eIF2B versus NSs-bound eIF2B after 3 hours, (**b**) apo eIF2B vs eIF2P-bound eIF2B after 3 hours, and (**c**) eIF2Bβγδε tetramer vs eIF2B(αβγδε)_2_ decamer after 100s. Positive values represent peptides with more protection (less deuteration) in the second listed state relative to the first listed state. Regions of NSs and eIF2-P protection on eIF2Bα are shown in orange and purple boxes. The eIF2Bδ C-terminal “Switch Helix” is shown in green boxes. eIF2P-dependent protection of eIF2Bγ and eIF2Bε are indicated by grey and red boxes, respectively. eIF2Bβ protection upon decamerization is indicated by the pink box. (**d-f**) Structural maps of: (**d**) the NSs binding pocket, (**e**) eIF2αP binding pocket, and (**f**) the tetramer-tetramer interface of the eIF2B(αβγδε)_2_ decamer. Regions of interest are color-coded, corresponding to protected regions shown in deuteration difference plots (see **a-c**) and representative peptide uptake plots (see **g-h**). (**g-h**) Peptide uptake plots, showing the total number of exchanged deuterons per condition over time for representative peptides (representing an average of three independent experiments; error bars S.E.M.).

A summary of all HDX-MS data is shown in Figure 3. Differential deuteration uptake of peptides across all regions of eIF2B are plotted (***Fig. 3a-c***), peptides showing increased protection (relative to the apo state) are mapped onto the corresponding cryo-EM structure (***Fig. 3d-f***), and representative uptake profiles show individual peptide deuteration over time (***Fig. 3g,h***). First, we looked at peptides corresponding to regions known to be involved in interactions between modulators and eIF2B. HDX-MS faithfully identified interaction interfaces identified in published Cryo-EM structures. For instance, comparing apo eIF2B to the NSs-stabilized A-State eIF2B, we observed increased protection for peptides in eIF2Bα that map to the known NSs binding interface (***Fig. 3a,d,g***; orange and purple indicate regions of NSs binding to eIF2Bα; ***Supp. Fig. 3***). We also observed protection in peptides at the 2BAct binding site (a groove between eIF2Bβ and eIF2Bδ’ on the opposite face of eIF2B) for the 2BAct-stabilized eIF2B A-State sample, but not for peptides at the NSs binding interface (***Supp. Fig. 5a*,** blue boxes indicate regions of 2BAct binding to eIF2B).

Similarly, we observed protection of known eIF2-P binding interfaces when comparing apo-eIF2B to the eIF2P-induced eIF2B I-State. eIF2αP binds in the same pocket targeted by NSs but forms additional contacts with the eIF2Bδ subunit to stabilize the I-state (***Fig. 3b,e,g***; orange and purple indicate regions of eIF2-P binding to eIF2Bα; ***Supp. Fig. 4***). The extended eIF2-P trimer also forms contacts and regions of increased protection on the eIF2Bγ subunit, which are consistent with structural models (***Fig. 3b***; grey boxes indicate regions of eIF2-P binding to eIF2Bγ). (We note that eIF2-P also increased protection in eIF2Bε peptides, specifically towards its C-terminal HEAT domain, which is not resolved in current cryo-EM structures (***Fig. 3b***; see red box)).

Excitingly, we also observed novel regions of protection not corresponding to modulators’ direct binding interfaces, which therefore must be a result of allosteric modulation. We observed an entirely unexpected protection in the NSs-bound A-State mapping to the most C-terminal eIF2Bδ helix (***Fig. 3a,d,g***; indicated in green). In the A-State stabilized by 2BAct we saw the same effect on the C-terminal eIF2Bδ helix albeit to a slightly lower extent (***Supp. Fig. 5a,b***). Importantly, this region does not show increased protection in the eIF2-P-bound I-State (***Fig. 3b,e,g***). These results are surprising because the C-terminal eIF2Bδ helix does not make direct contact with NSs or 2BAct (***Fig. 3d***). The fact that we see protection of the C-terminal eIF2Bδ helix in both of the A-State stabilized conditions suggests that biasing eIF2B into the A-State causes protection of this region allosterically rather than by direct contact with ligands.

### eIF2Bδ C-terminal helix also becomes protected in eIF2B’s tetramer→decamer transition

We next asked what local structural rearrangements mediate assembly of the eIF2B decamer by comparing the HDX-MS profiles of the eIF2B tetramer and decamer. We monitored deuteration of eIF2B tetramers alone versus those mixed with eIF2Bα_2_ dimers to form the decamer. Remarkably, peptides in the same eIF2Bδ C-terminal helix region emerged among the most differentially protected regions in the decamer (***Fig 3c,f,h***; indicated in green). Furthermore, the C-terminal helix of eIF2Bβ, which abuts the C-terminal eIF2Bδ helix across the trans-tetramer interface, also showed increased protection (***Fig 3c,f,h***; indicated in pink). Notably both of these regions of notably increased protection relative to other peptides located at the decamerization interface, which show a more moderate increase in protection.

Collectively, these data indicate that changes in the conformation of the eIF2Bδ C-terminal helix occur in both the A→I-State and the tetramer→decamer eIF2B conformational transitions. In the tetramer, the amide protons in this helix exchange rapidly, suggesting that it is poorly folded (***Fig. 3h***). Upon decamerization (to the apo-state decamer), these amide protons show increased protection on the timescale of the HDX experiment. Relative to the apo-State decamer, we see more rapid protection of the eIF2Bδ C-terminal helix in the A-state decamer (***Fig. 3g***). These data indicate that the eIF2Bδ C-terminal helix undergoes conformational and energetic changes in both assembly and A-State to I-State transitions. The fact that this helix is uniquely remodeled in both transitions suggests that it may play a crucial role in eIF2B activity regulation. Therefore, we probed the ability of specific molecular interactions with the helix’s sidechains to control the activity, conformation, and assembly of eIF2B.

### Defining the eIF2B Switch-Helix: functional implications of key sidechain interactions

We first asked how this helix participates in the eIF2B A→I-State transition. Then, we investigated its role in the tetramer→decamer transition, as discussed below. The eIF2Bδ C-terminal helix sequence is highly conserved (***Supp. Fig. 7***). Comparison of A- and I-State structures reveals that key side-chain interactions are profoundly remodeled (***Fig. 4a and Supp. Fig. 8***)^18,21^. In the A-State eIF2-eIF2B structure, eIF2B δR517 (δR517) forms a salt-bridge (salt-bridge distance 2.95 Å heavy-atom distance) with eIF2B αD298 (αD298) (PDB: 6O81). Upon transition to the I-State in response to eIF2-P binding to eIF2B, δR517 rotates by 60°, breaks its salt-bridge with αD298, and forms an alternate salt-bridge (salt-bridge distance 2.65 Å) with eIF2B δE445 (δE445) (Step 1 in ***Fig. 4b***) (PDB 6O9Z). Concomitantly, eIF2B δL516 (δL516) rotates 60°, and eIF2B δF443 (δF443) flips to an alternate rotamer position, avoiding steric clashes (Step 2 in ***Fig. 4b***). The side-change arrangement we see in the A-State are observed in all A-State structures, including the viral NSs-bound structure (PDB 7RLO), and the ISRIB-bound structure (7L7Z) (***Supp. Fig. 8***), suggesting that these arrangements reflect a general feature of the A→I-State transition.

**Fig. 4.**
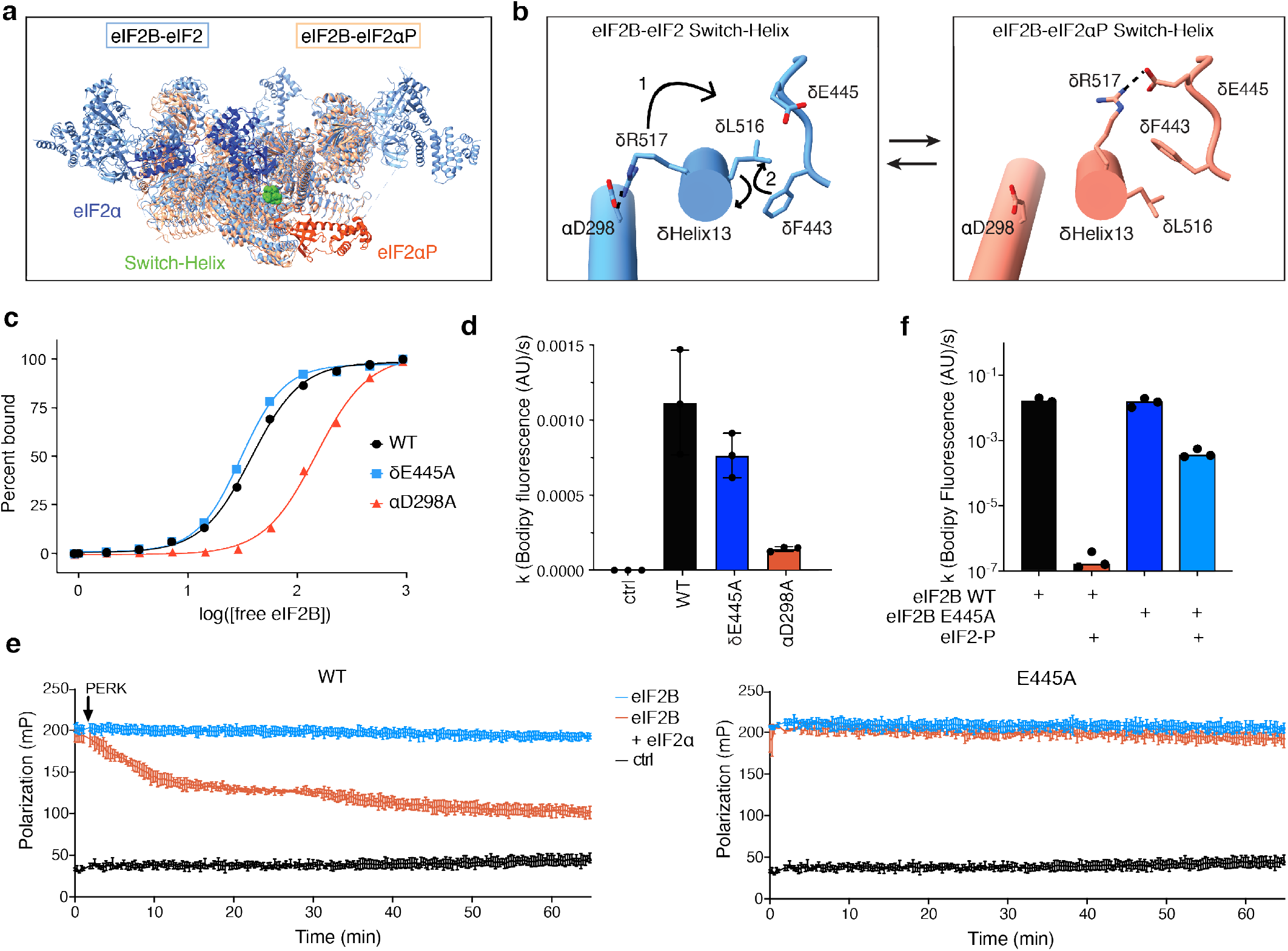
The eIF2Bδ C-terminal helix is a conformational switch. (**a**) Overview of atomic models of eIF2-bound eIF2B (light blue eIF2B, dark blue eIF2α, PDB 6o81) and eIF2P-bound eIF2B (salmon eIF2B, dark orange eIF2αP, PDB 6o9z) with the eIF2B C-terminal “Switch Helix” indicated in green. (**b**) The eIF2Bδ C-terminal helix undergoes a conformational change mediated by remodeled side-chain interactions in the A→I-state transition. In the A-state (left, PDB 6o81), eIF2Bδ residue R517 forms a salt bridge (dotted lines) with eIF2Bα D298 and the eIF2Bδ F443 rotamer is in the “down” position. (1) In the I-state (right, PDB 6o9z), eIF2Bδ R517 forms a new salt bridge (dotted lines) with eIF2Bδ E445, which coincides with (2) rotation of the eIF2Bδ C-terminal helix and adoption of the eIF2Bδ F443 “up” rotameric state. (**c**) Binding assay for fluorescent FAM-ISRIB interaction with eIF2B(αβγδε)_2_ decamers with indicated mutations using fluorescence polarization. Calculated EC_50_ values (95% confidence interval): WT=38±2 nM; δE445A=30 ±2 nM; D298A=152±18 nM. (**d**) Bodipy-GDP nucleotide loading assay ofeIF2B(αβγδε)_2_ decamers (final concentration 5 nM) with the indicated point mutation. Shown are averages and S.E.M. for three experimental replicates. (**e**) Kinetic fluorescence polarization dissociation assay for fluorescent FAM-ISRIB pre-indubated witheIF2B(αβγδε)_2_ decamers. At time zero, PERK kinase domain was spiked into the assay. Shown are averages and S.E.M. for three experimental replicates. (**f**) Bodipy-GDP nucleotide unloading assay of eIF2B(αβγδε)_2_ decamers (final concentration 5 nM) with or without point mutation and with or without the addition of 25 nM eIF2P, as indicated. Shown are averages and S.E.M. of rate constants (k) derived from a single exponential fit for three experimental replicates.

We posit that rotation of δR517 and δL516 and rearrangement of the salt-bridge and phenylalanine rotamer positions forms a concerted functional switch, and hence refer to the eIF2Bδ C-terminal helix as the “Switch-Helix”. According to this view, triggering this switch is required for the A→I-State transition, with the Switch-Helix facilitating allosteric communication between the inhibitor-binding site and other functional regions of eIF2B, such as the ISRIB- and/or the eIF2 substrate-binding pockets. To test this notion, we sought to bias the Switch-Helix toward a single conformation.

We first mutated eIF2B αD298 to alanine (αD298A) to promote the Switch-Helix I-State by aiming to prevent δR517 from engaging in the A-State salt-bridge interaction. Indeed, eIF2B αD298A displayed decreased binding of fluorescently-labeled ISRIB (FAM-ISRIB), as observed by fluorescence polarization (***Fig. 4c***) and previously reported for the I-State^14,22^. This suggests that the αD298A mutation allosterically affects the ISRIB binding pocket and induces the eIF2B I-State. Consistent with this interpretation, eIF2B αD298A also exhibited profoundly reduced nucleotide exchange activity (***Fig. 4d***). Using the same logic, we mutated eIF2B δE445 to alanine (δE445A), thus removing the I-State saltbridge and converting the Switch-Helix into the A-state. Neither FAM-ISRIB binding nor nucleotide exchange activity were diminished (***Fig. 4c,d***), consistent with an A-State phenotype that is assumed by apo-eIF2B^14^.

Based on these results, we predicted that converting the Switch-Helix into the A-State would render eIF2B more resistant to inhibition. To test this notion, we assessed FAM-ISRIB binding in response to the addition eIF2α-P. Indeed, binding of FAM-ISRIB to the eIF2B δE445A mutant was insensitive to inhibition by eIF2α-P (***Fig. 4e***). Similarly, eIF2B δE445A nucleotide exchange activity was less sensitive to inhibition by eIF2-P than eIF2B WT (***Fig. 4f***). These results are consistent with the interpretation that the δR517-δE445 salt-bridge plays an important role in stabilizing the I-State. In its absence eIF2B δE445A becomes biased toward the A-State, rendering eIF2-P inhibition less effective.

### Defining the eIF2B Switch-Helix: δF443 Phenylalanine rotamer

The second conformational element of the Switch-Helix suggested from the structural data concerns the interactions and rotameric disposition of eIF2B δF443 and δL516. eIF2B δF443 forms the “up” rotamer position in the I-State and the “down” rotamer position in the A-State, corresponding to a mirrored repositioning of δL516 (***Fig. 4d***). To explore the functional consequences of this rearrangement we mutated eIF2B δL516 and δF443 independently to alanine.

We first asked whether mutating these amino acids caused allosteric remodeling of the ISRIB pocket. Strikingly, we observed that the eIF2B δL516A mutation resulted in a decrease in FAM-ISRIB binding by almost an order of magnitude (8-fold), while the δF443A mutation caused reduced FAM-ISRIB binding by 2-fold (***Fig. 5a***). Similarly, eIF2B δL516A mutation strongly decreased nucleotide exchange activity, while eIF2B δF443A’s activity was only slightly reduced (***Fig. 5b***). These results are consistent with the interpretation that in the absence of the steric clash driven by δL516, the δF443 rotamer and eIF2B as a whole default to the I-state orientation. If there is no δF443 rotamer clash (such as in the eIF2B F443A mutant), the switch is broken and eIF2B can assume either A-State or I-State.

**Fig. 5.**
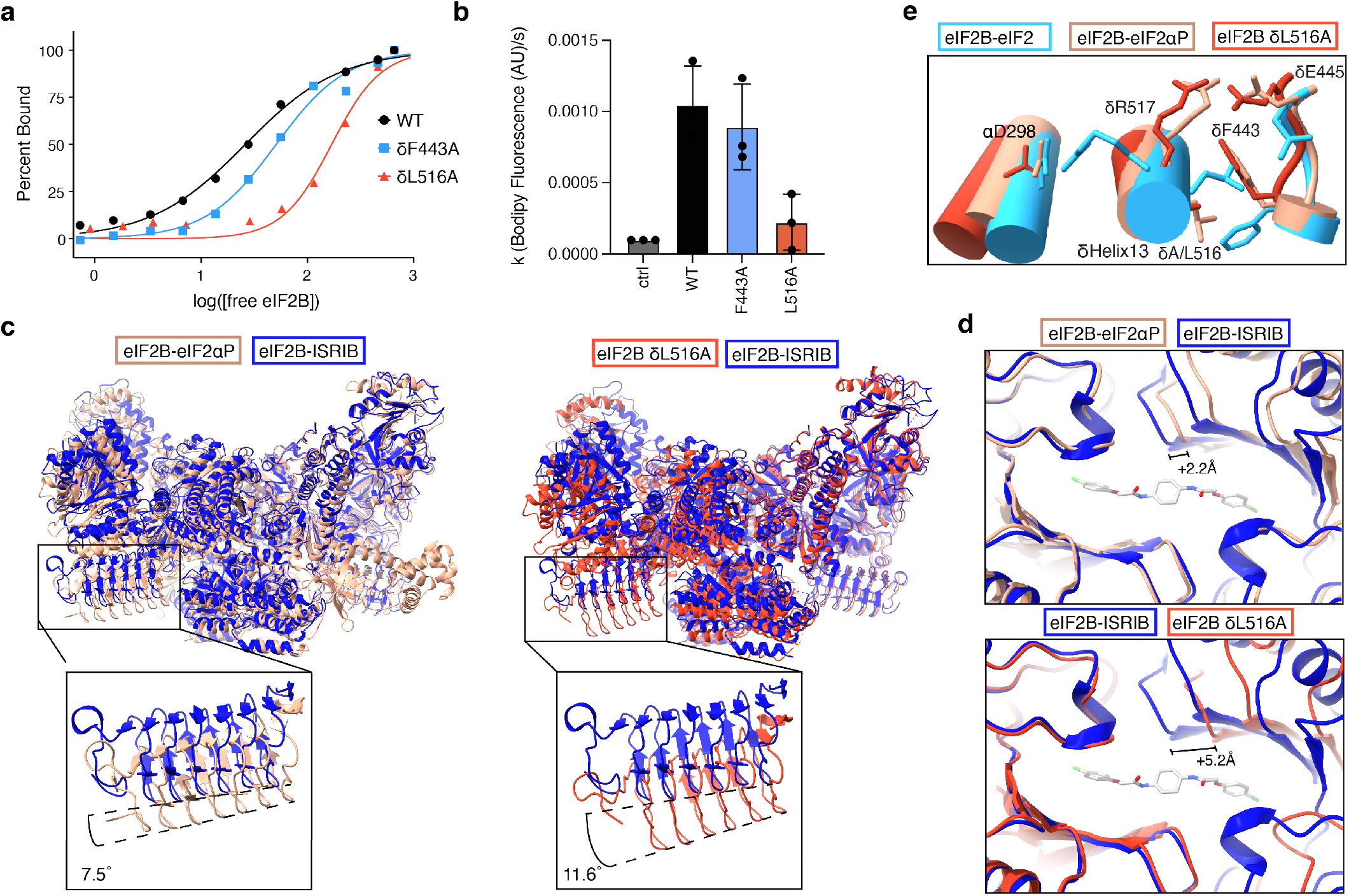
eIF2B Switch-Helix controls the A-to-I-State transition. (**a**) Binding assay for fluorescent FAM-ISRIB interaction with eIF2B(αβγδε)_2_ decamers containing the indicated mutations using fluorescence polarization. Calculated EC_50_ values (95% confidence interval): WT=26±3 nM; F443A=50 ±8 nM;δL516A=169±34 nM. (**b**) Bodipy-GDP nucleotide loading assay of eIF2B(αβγδε)_2_ decamers (final concentration 5 nM) with the indicated point mutation. Shown are averages and S.E.M. of rate constants (k) derived from a single exponential fit for three experimental replicates. (c) Atomic model of δL516A decamer structure (dark orange) overlaid on ISRIB-bound A-state eIF2B model (PDB 7L7G, blue) and eIF2αP-bound I-state eIF2B model (PDB 6o9z, peach). (**d**) Zoom-in view of the ISRIB binding pocket (PDB 7L7G), showing widening upon eIF2αP binding (PDB 6o9z, peach) and further widening for the δL516A decamer (dark orange). The 2.2 A° and 5.2 A° pocket lengthening was measured between eIF2Bβ Asn162 and eIF2B δ Ser178. (**e**) Overlay of δL516A Switch-Helix side chains (dark orange) onto eIF2-bound eIF2B A-state decamer (PDB 6o81, blue) and eIF2αP-bound eIF2B I-state (PDB 6o9z, peach) atomic models.

To further test this prediction, we determined a cryo-EM structure of eIF2B δL516A. After 2D and 3D classification, we generated a single consensus structure of the eIF2B δL516A decamer at 2.9 Å resolution (***Table 1, Supp. Fig. 9***) with most sidechain density clearly visible. We then built the atomic model of eIF2B δL516A into this map. Consistent with our predictions, eIF2B δL516A exhibited a “wings-down” I-State-like conformation. The two tetramer subcomplexes undergo a rocking motion that changes the angle between them by 11.6° relative to the ISRIB-bound A-state (***Fig. 5c***). Indeed, eIF2B δL516A adopted a more extreme I-State conformation than either the previously reported eIF2α-P-bound eIF2B (where tetrameric subcomplexes rotated by 7.5°) or the I-state-like eIF2B βH160D mutant (where tetrameric subcomplexes rotated by only 3.5°)^18,32^. Relative to ISRIB-bound A-State eIF2B^21^, the eIF2B δL516A ISRIB pocket widened by 5.2 Å, and the eIF2B-eIF2αP widened by 2.2 Å (**Fig. 5d**). Similarly, compared to the eIF2B-eIF2 structure^18^, the substrate binding pocket widened by 4.5 Å in the eIF2B δL516A structure, and 2.1 Å in the eIF2B-eIF2αP structure (***Supp. Fig 10*.**). Close examination of the eIF2B δL516A Switch-Helix confirmed that δF443 defaulted to an I-State rotameric position (***Fig. 5e and Supp. Fig. 8d***). Critically, in the δL516A structure, δR517 also formed the I-State salt-bridged conformation, reinforcing the notion that the salt-bridge and rotamer elements of the Switch-Helix function as a concerted switch.

**Table 1.** Cryo-EM data collection, analysis and model building.

| Structure | eIF2B<sup>δL516A</sup><br>(PDB ID: XXXX) | eIF2Bβγδε<br>(PDB ID: XXXX) |
| --- | --- | --- |
| **Data collection** |  |  |
| Microscope | Titan Krios |  |
| Voltage (keV) | 300 |  |
| Nominal magnification | 105000x |  |
| Exposure navigation | Image shift |  |
| Electron dose (e<sup>-</sup>Å<sup>-2</sup>) | 67 |  |
| Dose rate (e<sup>-</sup>/pixel/sec) | 8 |  |
| Detector | K3 summit |  |
| Pixel size (Å) | 0.835 |  |
| Defocus range (μm) | 0.6-2.0 |  |
| Micrographs | 4042 | 2692 |
| **Reconstruction** |  |  |
| Total extracted particles (no.) | 2729647 | 464356 |
| Final particles (no.) | 326478 | 71704 |
| Symmetry imposed | C1 | C1 |
| FSC average resolution, masked (Å) | 2.9 | 3.1 |
| FSC average resolution, unmasked (Å) | 3.7 | 3.9 |
| Applied B-factor (Å) | 89.5 | 115.4 |
| Reconstruction package | Cryosparc v3.3.2 | Relion 3 and Cryosparc v3.3.2 |
| **Refinement** |  |  |
| Protein residues | 3136 | 1325 |
| Ligands | 0 | 0 |
| RMSD Bond lengths (Å) | 0.008 | 0.004 |
| RMSD Bond angles (°) | 0.851 | 0.764 |
| Ramachandran outliers (%) | 0.06 | 0.31 |
| Ramachandran allowed (%) | 7.76 | 6.46 |
| Ramachandran favored (%) | 92.18 | 93.24 |
| Poor rotamers (%) | 11.91 | 10.98 |
| CaBLAM outliers (%) | 2.65 | 3.29 |
| Molprobity score | 2.79 | 2.70 |
| Clash score (all atoms) | 9.32 | 8.85 |
| B-factors (protein) | 85.13 | 54.09 |
| B-factors (ligands) | N/A | N/A |
| EMRinger Score | 2.90 | 3.45 |
| Refinement package | Phenix 1.17.1-3660-000 | Phenix 1.17.1-3660-000 |

To exclude the possibility that the mutations analyzed here impacted eIF2B decamerization, we analyzed mutant eIF2B complexes by sedimentation velocity analytical ultracentrifugation. We observed no defect in decamerization for eIF2B αD298A or δE445A and only minor defects for δL516A and δF443A (***Supp. Fig. 11***). These latter defects could be corrected by slightly increasing the concentration of eIF2Bα_2_, both in AUC experiments and in nucleotide exchange assays, indicating that this particular disruption of the Switch-Helix also slightly lowered the affinity for eIF2Bα_2_ (Supp Fig. 9).

### eIF2B Switch-Helix elements conformationally rearrange upon decamer assembly

We next returned to the HDX-MS observation of a selective increase in protection upon tetramer→decamer assembly for the peptides in the Switch-Helix (***Fig. 2c***). This result is notable, as the interface that becomes buried during decamer assembly includes many other structural elements for which no similar increase in protection was observed. This suggests that increased in protection is a result of local changes in the helix.

To explore conformational change and side-chain interactions in this region of increased protection associated with the assembly reaction, we determined a cryo-EM structure of eIF2Bβγδε tetramer to 3.1 Å resolution (***Fig. 6a, Table 1, Supp Fig. 12***). Overall, the architecture of the tetramer alone is very similar to that of the eIF2Bβγδε tetramer in the context of the eIF2B(αβγδε)_2_ decamer (***Fig. 6b***). A close look at the Switch-Helix elements revealed that the side chains in the eIF2Bβγδε tetramer adopt the “I-state” conformation, with δR517 forming a salt bridge with δE445 and the δF443 rotamer in the “up” position (***Fig. 6c, Supp Fig. 8d***), consistent with the increased protection seen for this region in HDX-MS.

**Figure 6:**
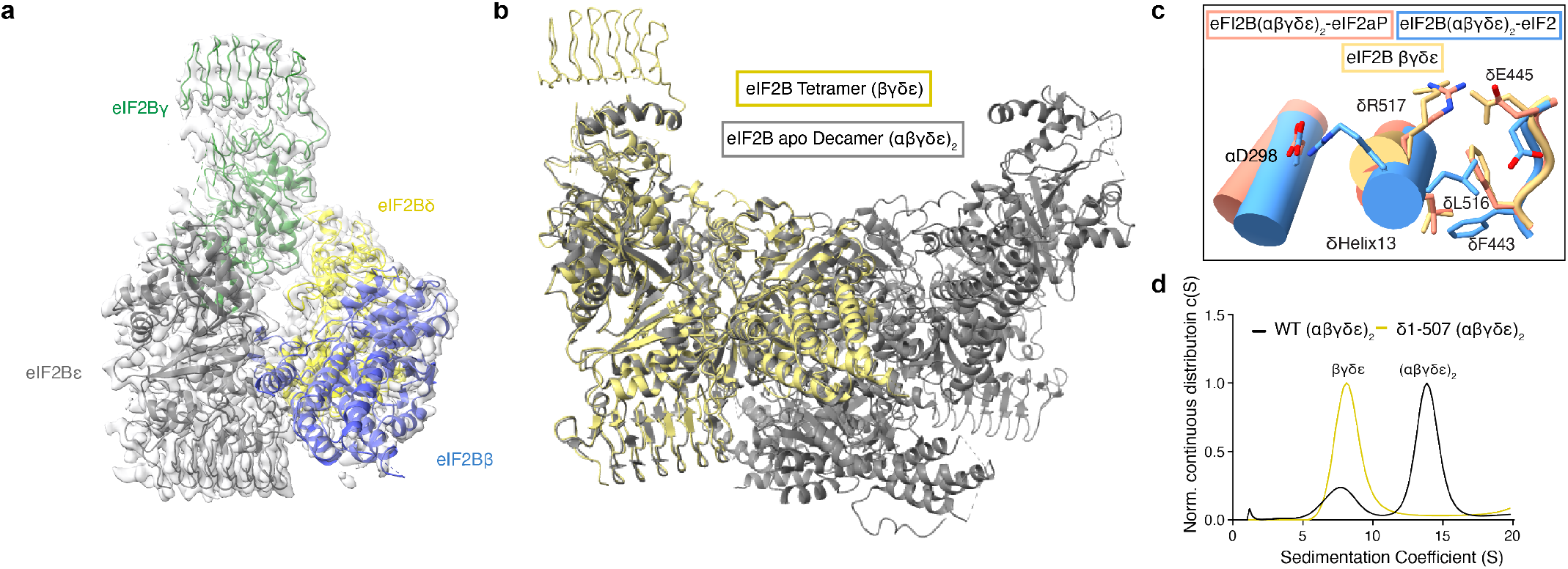
eIF2B Switch-Helix is triggered in the tetramer-to-decamer transition. (a) Atomic model of eIF2B tetramer overlaid with EM density. (b) Overlay of eIF2Bβγδε tetramer structural model onto apo eIF2B(αβγδε)_2_ decamer structural model (PDB 7L70). (c) Overlay of eIF2Bβγδε tetramer Switch-Helix side chains (yellow) onto eIF2-bound eIF2B A-state decamer (blue, PDB 6o81) and eIF2αP-bound eIF2B I-state (salmon, PDB 6o9z) atomic models. (d) Sedimentation velocity analytical ultracentrifugation analysis of WT eIF2B(αβγδε)_2_ and eIF2B(αβγδε)_2_ without the eIF2Bδ C-terminal helix (δ1-507).

To test the role of the Switch-Helix in decamerization, we performed analytical ultracentrifugation experiments on eIF2B truncated for δHelix13 (δ1-507αβγδε) and found that the Switch-Helix is required for decamerization (***Fig. 6d***). Hence, we conclude that protection of the Switch-Helix upon addition of the eIF2Bα_2_ dimer reflects both decamerization and the adoption of the A-state Switch-Helix conformation.

### Endogenous editing of Switch-Helix mutations into cells tunes ISR signaling

To determine how Switch-Helix orientation impacts ISR signaling *in cellula*, we introduced Switch-Helix mutations at the endogenous locus of pseudohaploid AN3-12 mouse ES cells, chosen to facilitate ease of editing. Using CRISPR/Cas9 technology, we obtained three homozygous eIF2Bδ E446A (homologous to human eIF2Bδ E445A) clones and two heterozygous eIF2Bα D298A/WT clones, as validated by sequencing. Importantly, eIF2Bα and eIF2Bδ protein levels remained comparable to the unedited parental cells, demonstrating that the mutation does not destabilize edited subunits or disrupt stoichiometry between eIF2B complex members (***Fig. 7a,b***). Consistent with predictions for A-State stabilization, eIF2Bδ E446A homozygous clones showed reduced ATF4 induction upon treatment with thapsigargin (Tg), a small molecule inhibitor of the ER-Ca^++^ pump and potent inducer of the ISR (***Fig. 7a,c***). Consistent with expectations of I-State stabilization, eIF2Bα D298A heterozygous clones showed increased induction of ATF4 upon Tg treatment (***Fig. 7b,c***). We did not recover any homozygous αD298A clones, likely because the extreme I-state caused by this mutation induces strong ISR signaling that limits cellular proliferation. However, the observation of increased ATF4 induction upon heterozygous introduction of the D298A mutation is consistent with the interpretation that cells bearing a mixture of WT and D298A eIF2Bα are primed to launch a stronger ISR response than unedited cells (***Fig. 7b,c***).

**Figure 7:**
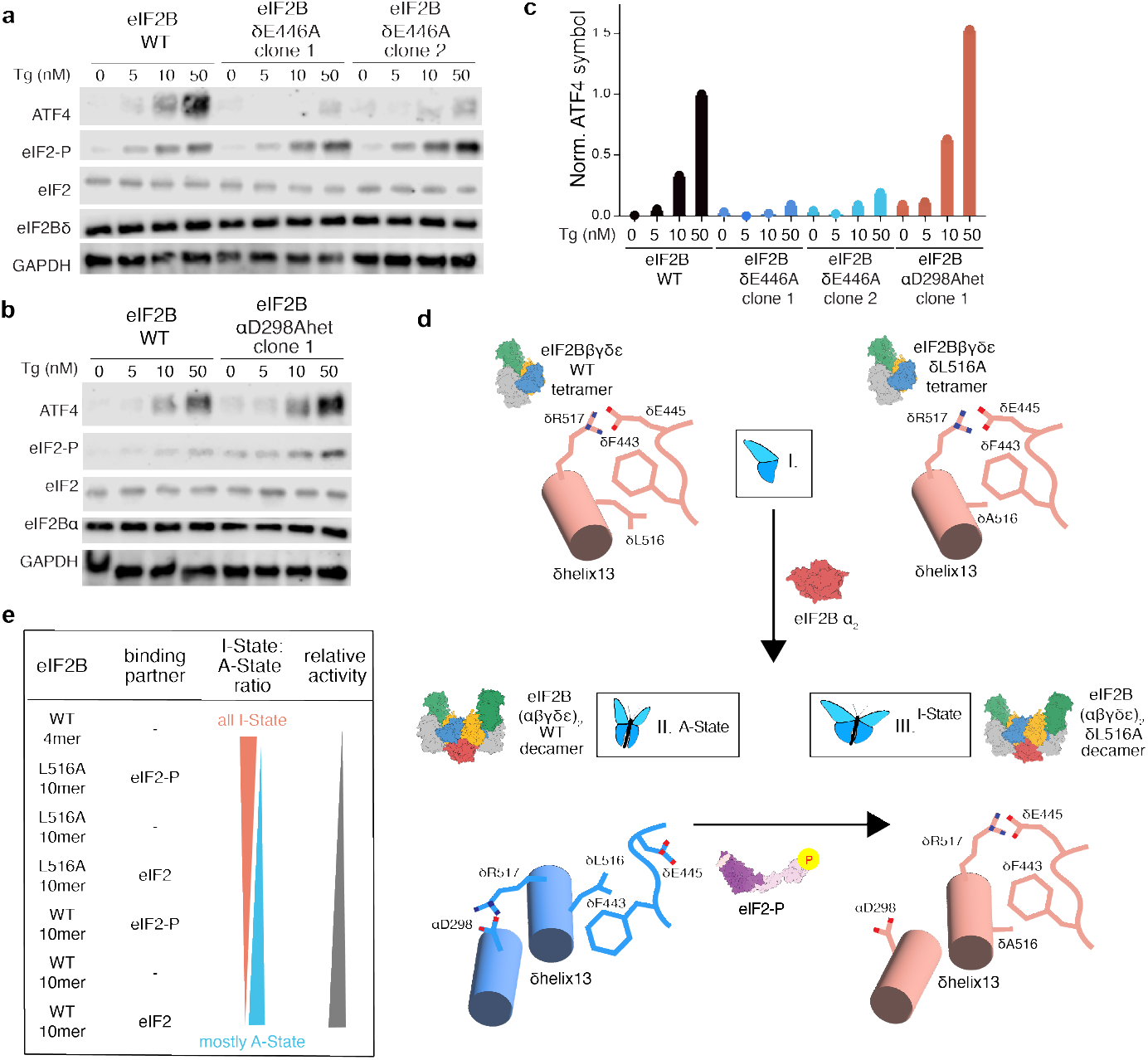
eIF2B Switch-Helix mutations control ISR signaling in cells. (**a, b**) AN3-12 mouse ES cells containing (**a**) eIF2BδE446A homozygous or (**b**) eIF2BαD298A heterozygous endogenously edited mutations were treated with the indicate concentration of Tg for 1 hour and immunoblotted for the indicated proteins. (**c**) Quantitation of ATF4 immunoblot intensity for blots shown in a,b. Signal was normalized to WT AN3-12 50 nM Tg ATF4 signal. (**d**) I. eIF2B Switch-Helix is in the I-State orientation in eIF2B βγδε tetramer. II. Incorporation of eIF2Bα_2_ prompts formation of the δR517-αD298 salt bridge causing a δL516-δF443 steric clash, triggering conformational change of the Switch-Helix and global conversion to the eIF2B A-State. III. For the eIF2B L516A variant, in the absence of the δL516-δF443 steric clash, the Switch-Helix is not triggered and global I-state conformation is maintained after decamerization. Finally, addition of eIF2-P also converts the Switch-Helix from the A-State to the I-State, reverting side-chains back to the same “I-State” arrangement assumed in the tetramer. (**e**) Model: eIF2B is in constant equilibrium between the I-State and A-State. I-State:A-State ratio is altered by Switch-Helix mutation and binding partner.

## Discussion

Through comprehensive characterization of eIF2B remodeling events via HDX-MS, Cryo-EM, biochemistry, and signaling assays, we discovered that allosteric coordination is mediated by a highly conserved switch that is triggered both in A→I-State and tetramer→decamer transitions. This Switch-Helix is located at the core of the eIF2B decamer and facilitates the hinging motion that converts eIF2B from the A-State to the I-State (***Figs. 1a and 4a***). Furthermore, the Switch-Helix is located between the eIF2 and eIF2αP binding pockets, enabling substrate and inhibitor-binding to engage the switch via local interactions (***Fig. 4a***). Remarkably, the Switch-Helix can also be triggered allosterically by binding of 2BAct or ISRIB to a pocket located 40Å away, showing that it is part of an inter-subunit allosteric communication network that couples substrate-binding, inhibitor-binding, and ISRIB-binding pockets (***Supp. Movie 1***).

A concerted conformational change in the Switch-Helix drives both eIF2B’s assembly and A-State to I-State transitions. To summarize, our results indicate that Switch-Helix orientation is identical in the eIF2B tetramer and in the I-State eIF2B decamer, and adopts a different state in the A-State decamer. Helix conformation therefore correlates with eIF2B activity. We pose a model in which concerted interactions between side-chain rotamers are responsible for this switch. In the context of the isolated eIF2B tetramer, the δF443 rotamer is in the “up” position and δR517 is salt-bridged to δE445 (***Fig. 7d*** step I). In this rotameric state, the δF443 sidechain interacts with the sidechain from δL516, which is adjacent to the δR517-δE445 salt bridge. In contrast, upon decamerization to the A-State, these same residues are involved in different interactions: δR517 now forms a salt bridge to αD298, a residue not present in the tetramer (***Fig. 7d***, step II). In the presence of this alternative salt-bridge, L516 is forced into a position that would clash with the “up” F443 rotameric state. Thus, through these concerted changes, formation of the decamer forces the F443 rotamer to adopt the “down” A-State conformation.

Binding of eIF2-P inhibitor to the A-State decamer results in the eIF2B I-State, and induces the same concerted shift, restoring the δF443 “Up” rotamer (***Fig. 7d***, step III). We propose that the lynchpin in this mechanism is steric restriction on the L516 sidechain. In the absence of the leucine sidechain (such as in the δL516A variant) these restrictions are released and δF443 is able to adopt its intrinsically lowest-energy conformation (the “up” I-State) (***Fig. 7d***, step III). Thus, this domino of effects of local side-chain interactions involving a single central helix define an allosteric mechanism for activation of the entire eIF2B complex. These local changes in the helix are propagated to effector binding regions via tuning the relative orientation between eIF2Bα and eIF2Bδ subunits. Hence, the Switch-Helix acts as a fulcrum to tune ISR signaling.

These small local Switch-Helix changes are easily promoted by the binding energies and interfaces of eIF2Bα_2_ or allosteric modulators such as eIF2 substrate or eIF2-P inhibitor. Notably, binding of eIF2-P inhibitor also leverages coupled interactions within the Switch-Helix fulcrum by providing sufficient binding energy to flip the Switch-helix into the I-state (***Fig. 7d***). It is fascinating to consider whether the Switch-Helix preceded or co-evolved with eIF2-P inhibitor function. Apo-eIF2B has the capacity to adopt an A-State or I-State orientation, as demonstrated by the I-state orientation of the tetramer switch-helix and by the fact that apo-eIF2B L516A adopts the I-State. This suggests that eIF2-P binding licenses the intrinsic capacity of the eIF2B Switch-helix to induce to the I-State. Overall, the fact that a shared set of amino acid residues centrally participate in both conformational and assembly eIF2B regulatory transitions highlights that the Switch-Helix is the core allosteric node regulating eIF2B state.

Our model of allosteric activation rests on the ability of eIF2B to conformationally sample two discrete states (***Fig. 7e***). Covariation analysis performed on apo eIF2B cryo-EM datasets shows that the population of single particles samples both the A-State and the I-State, but on average assume the A-State^32^. This explains why the apo-eIF2B HDX-MS profile looks similar to the I-state eIF2B-eIF2-P HDX-MS profile. Assuming the dynamic sampling of the I- and A-State occurs on a time-scale faster than the HDX-MS experimental timescale, the HDX-MS profile would be dominated by the faster exchanging species (the I-State), even if it was not the majority of the population.

ISR signaling outcomes in the cell depend on the relative population of A-State and I-State eIF2B molecules, which can be biased by binding partners such as eIF2 substrate and eIF2-P inhibitor. Switch-Helix side-chain interactions define these relative populations of A-State and I-State eIF2B. Notably we observe that Switch-Helix mutations tune activity without completely impeding conformational change induced by external binding partners such as eIF2-P (*see **Fig. 4f***). As such, the Switch-Helix is a metastable bistable switch; while it must adopt one of two discrete states, the system is tuned such that relevant biological interactions control the relative population of the two states. Importantly, the eIF2B δL516A structure forms a “super-I-State”, with a wider overall rotation across the central hinge than is observed in the eIF2-P bound structure. This suggests that δL516A more strongly shifts the population of eIF2B conformations toward the I-State than does eIF2-P.

The dynamic structural insights presented here suggest new strategies for development of smallmolecule therapies. Stabilization of the Switch-Helix into the A-State would be predicted to have ISRIB-like effects on ISR signaling in cells and could be accomplished by pharmacological targeting of the I-State specific δR517-δE445 salt bridge. Furthermore, we show that endogenously editing the δE446A mutation into cells programs in an ISRIB-like state, which opens numerous possibilities for new *in vivo* studies.

Overall, this work integrates multiple structural and biochemical techniques to identify the Switch-Helix as a central allosteric mechanism that controls both eIF2B conformational and assembly state transitions. Through small concerted changes this central Switch-Helix functions as a molecular fulcrum to affect signaling functions greater than 140Å away (in the case of the ISRIB binding pocket). Remarkably, HDX-MS allowed us to identify the single helix responsible for this allosteric transition. Further structural investigation using Cryo-EM identified side-chain interactions that act as highly conserved allosteric hotspots, shaping eIF2B activity and ISR signaling. Finally, by manipulating these sidechains via mutation, we were able to confirm effects of these subtle changes in Switch-Helix orientation on ISR signaling in cells. This atomic level understanding of eIF2B’s allosteric mechanism provides the needed foundation for rational control of ISR signaling and adds a new dimension to the molecular understanding of allosteric mechanisms in such large regulatory complexes.

## Materials and Methods

### Purification and assembly of human eIF2B subcomplexes

Human eIFBα2 (pJT075), eIF2Bβγδε (pJT073 and pJT074 co-expression), and eIF2Bβγδε-F (pMS029 and pJT074 co-expression) were purified as previously described^16^.

### Purification of viral NSs protein

Viral NSs::6xHIS was purified as previously described^26^. Briefly, we used the pMS113 construct to express and purify NSs::6xHIS. Expi293T cells (Thermo Fisher) were transfected with the NSs construct per the manufacturer’s instructions for the MaxTiter protocol and harvested 5 days after transfection. Cells were pelleted (1000g, 4 min) and resuspended in Lysis Buffer (130 mM KCl, 2.5 mM MgCl2, 25 mM HEPES-KOH pH 7.4, 2 mM EGTA, 1% triton, 1 mM TCEP, 1× cOmplete protease inhibitor cocktail (Roche)). Cells were then incubated for 30 min at 4 °C and then spun at 30,000g for 1 h to pellet cell debris. Lysate was applied to a 5 ml HisTrap HP column (GE Healthcare) equilibrated in Buffer A (20 mM HEPES-KOH, pH 7.5, 200mM KCl, 5 mM MgCl2, 15 mM imidazole) and then eluted using a gradient of Buffer B (20 mM HEPES-KOH, pH 7.5, 200 mM KCl, 5 mM MgCl2, 300 mM imidazole). NSs::6xHIS was concentrated using a 10 kDa MWCO spin concentrator (Amicon) and further purified by size exclusion chromatography over a Superdex 200 Increase 10/300 GL column (GE Healthcare) in Elution Buffer (20 mM HEPES, pH 7.5, 200 mM KCl, 5 mM MgCl2, 1mM TCEP, and 5% Glycerol). The resulting fractions were pooled and flash frozen in liquid nitrogen.

### Purification of heterotrimeric human eIF2

All experiments performed prior to March 1, 2022 were performed using human eIF2 purified as previously described^27^. Some experiments were performed using purified eIF2 that was a generous gift of Calico Life Sciences LLC. All experiments performed after March 1, 2022 used eIF2 purified according to a second previously described protocol^15^.

### Phosphorylation of eIF2 trimer (eIF2-P) and eIf2α (eIF2αP)

To generate phosphorylated eIF2 trimers or eIF2α, 25 μM eIF2 trimer or eIF2α were incubated with 500 nM recombinant PERK-kinase domain (purified in-house as previously described^14^) and 1 mM ATP at 37C for 1 hour. Phosphorylation of the final product was verified by 12.5% SuperSep PhosTag gel (Wako Chemical Corporation).

### Purification and assembly of human eIF2B subcomplexes

Human eIFBα2 (pJT075), eIF2Bβγδε (pJT073 and pJT074 co-expression), and eIF2Bβγδε-F (pMS029 and pJT074 co-expression) were purified as previously described^21^.

### Purification of viral NSs protein

Viral NSs::6xHIS was purified as previously described^32^. Briefly, we used the pMS113 construct to express and purify NSs::6xHIS. Expi293T cells (Thermo Fisher) were transfected with the NSs construct per the manufacturer’s instructions for the MaxTiter protocol and harvested 5 days after transfection. Cells were pelleted (1000g, 4 min) and resuspended in Lysis Buffer (130 mM KCl, 2.5 mM MgCl2, 25 mM HEPES-KOH pH 7.4, 2 mM EGTA, 1% triton, 1 mM TCEP, 1× cOmplete protease inhibitor cocktail (Roche)). Cells were then incubated for 30 min at 4 °C and then spun at 30,000g for 1 h to pellet cell debris. Lysate was applied to a 5 ml HisTrap HP column (GE Healthcare) equilibrated in Buffer A (20 mM HEPES-KOH, pH 7.5, 200mM KCl, 5 mM MgCl2, 15 mM imidazole) and then eluted using a gradient of Buffer B (20 mM HEPES-KOH, pH 7.5, 200 mM KCl, 5 mM MgCl2, 300 mM imidazole). NSs::6xHIS was concentrated using a 10 kDa MWCO spin concentrator (Amicon) and further purified by size exclusion chromatography over a Superdex 200 Increase 10/300 GL column (GE Healthcare) in Elution Buffer (20 mM HEPES, pH 7.5, 200 mM KCl, 5 mM MgCl2, 1mM TCEP, and 5% Glycerol). The resulting fractions were pooled and flash frozen in liquid nitrogen.

### Purification of heterotrimeric human eIF2

All experiments performed prior to March 1, 2022 were performed using human eIF2 purified as previously described^33^. Some experiments were performed using purified eIF2 that was a generous gift of Calico Life Sciences LLC. All experiments performed after March 1, 2022 used eIF2 purified according to a second previously described protocol^19^.

### Phosphorylation of eIF2 trimer (eIF2-P) and eIf2α (eIF2αP)

To generate phosphorylated eIF2 trimers or eIF2α, 25 μM eIF2 trimer or eIF2α were incubated with 500 nM recombinant PERK-kinase domain (purified in-house as previously described^18^) and 1 mM ATP at 37C for 1 hour. Phosphorylation of the final product was verified by 12.5% SuperSep PhosTag gel (Wako Chemical Corporation).

### Assembly of eIF2B decamer complexes

All eIF2B(αβγδε)_2_ used throughout was assembled by mixing purified eIF2Bβγδε and eIF2Bα_2_ at a molar ratio of 2:1.2 eIF2Bβγδε:eIF2Bα_2_ unless otherwise indicated. Complexes were assembled at 10 μM and incubated at 4C for 30 minutes prior to dilution to experimental conditions.

### Assembly of eIF2B complexes for HDX-MS characterization

For HDX-MS experiments, after preparation of 10 μM eIF2B(αβγδε)_2_, 10x mixtures containing 5 μM eIF2B (αβγδε)_2_ and 12 μM NSs or eIF2-P (representing a 1.2-fold molar excess of NSs and eIF2-P relative to available eIF2B binding sites) were prepared in HX buffer and allowed to assemble overnight at 4C. 10x 2BAct samples were prepared by combining 5 μM eIF2B with 10 μM 2BAct dissolved in DMSO. Matched 10x control samples were prepared by combining 5 μM eIF2B with 2% DMSO such that the final experimental DMSO concentration was 0.2%.

### Hydrogen Deuterium Exchange

For all hydrogen deuterium exchange experiments, deuterated buffer was prepared by lyophilizing eIF2B assay buffer (20 mM HEPES, 200 mM KCl, 5 mM MgCl_2_, 1 mM TCEP, pH 7.9) and resuspending it in D_2_O (Sigma-Aldrich 151882). To initiate the continuous-labeling experiment, samples were diluted tenfold to 1X (final eIF2B tetramer concentration of 1 μM or eIF2B decamer concentration of 0.5 μM) into temperature-equilibrated, deuterated eIF2B assay buffer. Samples were quenched at the time points outlined below, by mixing 30 μL of the partially exchanged protein with 30 μL of 2× quench buffer (6M Urea, 500 mM TCEP, pH 2.4) on ice. Samples were incubated on ice for 1 minute to allow for partial unfolding to assist with proteolytic degradation and then were flash frozen in liquid nitrogen and stored at −80 °C. The hydrogen deurterium exchange time points for these experiments were 10 seconds, 100 seconds, 15 minutes, and 3 hours.

### Protease digestion and LC–MS

All samples were thawed immediately before injection into a cooled valve system (Trajan LEAP) coupled to a LC (Thermo UltiMate 3000). Sample time points were injected in random order. The temperature of the valve chamber, trap column, and analytical column were maintained at 2 °C. The temperature of the protease column was maintained at 10 °C. The quenched sample was subjected to in-line digestion by two immobilized acid proteases in order, aspergillopepsin (Sigma-Aldrich P2143) and porcine pepsin (Sigma-Aldrich P6887) at a flow rate of 200 μL/minute of buffer A (0.1% formic acid). Protease columns were prepared in house by coupling protease to beads (Thermo Scientific POROS 20 Al aldehyde activated resin 1602906) and packed by hand into a column (2 mm ID × 2 cm, IDEX C-130B). Following digestion, peptides were desalted for 4 minutes on a hand-packed trap column (Thermo Scientific POROS R2 reversed-phase resin 1112906, 1 mm ID × 2 cm, IDEX C-128). Peptides were then separated with a C8 analytical column (Thermo Scientific BioBasic-8 5-μm particle size 0.5 mm ID × 50 mm 72205-050565) and a gradient of 5–40% buffer B (100% acetonitrile, 0.1% formic acid) at a flow rate of 40 μL/min over 14 minutes, and then of 40–90% buffer B over 30 seconds. The analytical and trap columns were then subjected to a sawtooth wash and equilibrated at 5% buffer B prior to the next injection. Protease columns were washed with two injections of 100 μL 1.6 M GdmCl, 0.1% formic acid prior to the next injection. Peptides were eluted directly into a Q Exactive Orbitrap Mass Spectrometer operating in positive mode (resolution 70000, AGC target 3 × 10^6^, maximum IT 50 ms, scan range 300–1,500 *m/z*). For each eIF2B condition, a tandem mass spectrometry experiment was performed (Full MS settings the same as above, dd-MS^2^ settings as follows: resolution 17,500, AGC target 2 × 10^5^, maximum IT 100 ms, loop count 10, isolation window 2.0 *m/z*, NCE 28, charge state 1 and ≥7 excluded, dynamic exclusion of 15 seconds) on undeuterated samples. LC and MS methods were run using Xcalibur 4.1 (Thermo Scientific).

### Peptide identification

Byonic (Protein Metrics) was used to identify peptides in the tandem mass spectrometry data. Sample digestion parameters were set to non-specific. Precursor mass tolerance and fragment mass tolerance was set to 6 and 10 ppm respectively. Peptide lists (sequence, charge state, and retention time) were exported from Byonic and imported into HDExaminer 3 (Sierra Analytics). When multiple peptide lists were obtained, all were imported and combined in HDExaminer 3.

### HDExaminer 3 analysis

Peptide isotope distributions at each exchange time point were fit in HDExaminer 3. Deuteration levels were determined by subtracting mass centroids of deuterated peptides from undeuterated peptides.

### Bodipy-GDP exchange assay

In vitro detection of GDP binding to eIF2 was adapted from a published protocol for a fluorescence intensity–based assay describing dissociation of eIF2 and nucleotide^34^. We first performed a loading assay for fluorescent BODIPY-FL-GDP as described^21^. Purified eIF2 (137.5 nM) was incubated with 100 nM BODIPY-FL-GDP (Thermo Fisher Scientific) in assay buffer (100 mM HEPES-KOH, pH 7.5, 100 mM KCl, 5 mM MgCl2, 1 mM TCEP, and 1 mg/ml BSA) to a volume of 500 μl in a black-walled 1.5 mL tube. This mix was then added to 384 square-well black-walled, clear-bottom polystyrene assay plates (Corning Product #3766), 18 μl per well. A GEF mix composed of 10× solution of eIF2B(αβγδε)_2_ was prepared. To compare nucleotide exchange rates, 2 μl of the 10× GEF mixes were spiked into the 384-well plate wells with a multichannel pipette, such that the resulting final concentration of eIF2B(αβγδε)_2_ was 5 nM, and the final concentration of other proteins and drugs are as indicated in the figures. Subsequently, in the same wells, we performed a ‘GDP unloading assay’ as indicated in the figures. After completion of the loading reaction, wells were next spiked with 1 mM GDP to start the unloading reaction at t = 0. In the case of inhibition assays with eIF2-P, the eIF2/Bodipy-GDP mix was also incubated with 25 nM eIF2B(αβγδε)_2_ and a 10x mix of eIF2-P was spiked into the wells so that the final concentration was 50 nM. For all GEF assays involving eIF2-P an ‘unloading’ assay was used since the eIF2B(αβγδε)_2_ had been preincubated with eIF2. Fluorescence intensity was recorded every 10 s for 60 min at 25°C using a Clariostar PLUS (BMG LabTech) plate reader (excitation wavelength: 497 nm, bandwidth 14 nm; emission wavelength: 525 nm, bandwidth: 30 nm). Data collected were fit to a first-order exponential.

### FAM-ISRIB binding assay

All fluorescence polarization measurements were performed in 20 μl reactions with 100 nM eIF2B(αβγδε)_2_ + 2.5 nM FAM-ISRIB (Praxis Bioresearch) in FP buffer (20 mM HEPES-KOH pH 7.5, 100 mM KCl, 5 mM MgCl2, 1 mM TCEP) and measured in 384-well non-stick black plates (Corning Product #3820) using the ClarioStar PLUS (BMG LabTech) at room temperature. Prior to reaction setup, eIF2B(αβγδε)_2_ was assembled in FP buffer using eIF2Bβγδε and eIF2Bα_2_ in 2:1.5 molar ratio for at least 1 hour at 4°C. FAM-ISRIB was always first diluted to 2.5 μM in 100% NMP prior to dilution to 50 nM in 2% NMP and then added to the reaction. For titrations with eIF2α and eIF2α-P, dilutions were made in FP buffer. For titrations with ISRIB, it was first diluted with 100% NMP to 2.5 μM and then to the final concentrations in 4% NMP. The reactions with eIF2B, FAM-ISRIB, and these dilutions were incubated at 25°C for 30 min prior to measurement of parallel and perpendicular intensities (excitation: 482 nm; emission: 530 nm).

### FAM-ISRIB kinetic binding assay

The kinetic characterization of FAM-ISRIB binding during eIF2α phosphorylation was assayed in 18 μl reactions of 100 nM eIF2B(αβγδε)_2_, 2.5 nM FAM-ISRIB, 100 μM ATP, and 5.6 μM eIF2α/eIF2α-P in FP buffer. These solutions were pre-incubated at 22°C for 30 min before polarization was measured every 15 s (30 flashes/s). After four cycles, 2 μl of homemade PERK-KinaseDomain at 0.1 mg/mL was added for a final concentration of 10 μg/mL in the reaction, and measurement was resumed for 1 hour.

### Analytical ultracentrifugation

Analytical ultracentrifugation sedimentation velocity experiments were performed as previously described^21^.

### Sample preparation for cryo-EM microscopy

Decameric eIF2Bδ^L516A^ was prepared by incubating 20 μM eIF2Bδ^L516A^ βγδε with 11 μM eIF2Bα_2_ in a final solution containing 20 mM HEPES-KOH, 200 mM KCl, 5 mM MgCl2, and 1 mM TCEP. eIF2B eIF2Bδ^L516A^ decamers and eIF2B tetramers were diluted to 750 nM in 20 mM HEPES-KOH, 200 mM KCl, 5 mM MgCl2, and 1 mM TCEP prior to grid application. For grid freezing, a 3 μl aliquot of the sample was applied onto the Quantifoil R1.2/1/3 400 mesh Gold grid followed by a 30s waiting period. A 0.5 μl aliquot of 0.1-0.2% Nonidet P-40 substitute was added immediately before blotting. The entire blotting procedure was performed using Vitrobot (FEI) at 10 °C and 100% humidity.

### Electron microscopy data collection

Cryo-EM data was collected on a Titan Krios transmission electron microscope operating at 300 keV. Micrographs were acquired using a Gatan K3 direct electron detector. The total dose was 67 e-/ Å^2^, and 117 frames were recorded during a 5.9 s exposure. Data was collected at 105,000 x nominal magnification (0.835 Å/pixel at the specimen level), with a nominal defocus range of −0.6 to −2.0 μm.

### Image processing

The micrograph frames were aligned using MotionCor2^35^. The contrast transfer function (CTF) parameters were estimated with GCTF^36^. For the decameric eIF2Bδ^L516A^, Particles were picked in Cryosparc v3.3.2 using the apo eIF2B (EMDB: 23209) as a template^14,37^. Particles were extracted using an 128-pixel box size and classified in 2D. Classes that showed clear protein features were selected and extracted for heterogeneous refinement using models of an apo decamer, a tetramer and an impurity class, followed by homogenous refinement. Particles belonging to the decamer class were then re-classified using heterogeneous refinement to sort of the best resolution class. Particles from the resulting best class was then re-extracted with a pixel size of 0.835 Å, and then subjected to nonuniform refinement, yielding a reconstruction of 3.0 Å. These particles were subjected to CTF refinement to correct for the per-particle CTF as well as beam tilt. A final round of nonuniform refinement yielded the final structure of 2.9 Å.

For the tetramer structure, particles were picked by Gautomatch and extracted at a pixel size of 3.34 Å/pixel. Particles were imported into Relion 3.0 for autorefinement to generate a consensus structure. These particles were then subjected to multiple rounds of 2D classification, where particles that represent proteins are selected and re-extracted. The resulting set of particles were subjected to autorefinement, followed by re-extraction at 1.67 Å/pixel, and another round of autorefinement, yielding a reconstruction at 4.5 Å. These particles were then subjected to 3D classification (k = 4), and the best class was selected for further refinement, which generated a 4.2 Å reconstruction. Particles belonging to this set (72 k) are extracted at 0.835 Å/pixel and subjected to autorefinement, yielding a 3.9 Å structure. Particles belonging to this class are imported into Cryosparc v3.3.2, where they are subjected to nonuniform refinement, and CTF refinement to yield the final structure at the resolution of 3.1 Å.

### Atomic model building, refinement, and visualization

For both the decamer and the tetramer structures, the previously published apo eIF2B model (PDB ID: 7L70) was used as a starting model^14^. Each subunit was docked into the EM density individually and then subjected to rigid body refinement in Phenix^38^. The models were then manually adjusted in Coot and then refined in phenix.real_space_refine using global minimization, secondary structure restraints, Ramachandran restraints, and local grid search^39^. Then iterative cycles of manual rebuilding in Coot and phenix.real_space_refine were performed. The final model statistics were tabulated using Molprobity^40^. Molecular graphics and analyses were performed with the UCSF Chimera package. UCSF Chimera is developed by the Resource for Biocomputing, Visualization, and Informatics and is supported by NIGMS P41-GM103311. The atomic model is deposited into PDB under the accession codes XXXX (eIF2Bδ^L516A^) and XXXX (tetramer). The EM map is deposited into EMDB under the accession codes EMD-XXXXX (eIF2Bδ^L516A^) and EMD-XXXXX (tetramer).

### Generation of endogenously-edited cells

Editing of the EIF2B1 and eIF2B4 genes to introduce D298A and E446A mutations, respectively, into mouse AN3-12 pseudohaploid ES cells obtained from the Austrian Haplobank (haplobank.org) was performed using nucleofection of CRISPR-Cas9 RNPs as previously described (Floor lab protocol: https://www.protocols.io/view/cas9-sgrna-ribonucleoprotein-nucleofection-using-l-261ge1xyv479/v10). Nucleofection was performed using a 4D-nucleofector with X-unit attachment (Lonza) and with pulse program CG-104. Two days post-nucleofection, genomic DNA was extracted using the PureLink Genomic DNA mini kit from a portion of cells, relevant genes were PCR-amplified, and editing efficiency was determined using the Synthego ICE tool (https://ice.synthego.com/#/). When editing efficiency was confirmed to be at least 10%, cells were diluted to an expected density of 0.0625 cells/well and plated onto 96-well plates, then allowed to grow up from single colonies, with media changes every 3-5 days depending on media acidification rate. Genomic DNA was extracted from colonies derived from single clones, and successful editing was determined by PCR amplification of the gene of interest and analysis using Synthego ICE tool. All cell lines were negative for mycoplasma contamination.

### Western blotting

Cells were seeded at 1E6 cells/well of a 6-well plate and grown at 37°C and 5% CO_2_ for 24 h. Cells were treated using 10 nM thapsigargin (Tg) (Invitrogen) for 1 h, ensuring the final DMSO concentration was 0.1% across all conditions. Plates were put on ice, cells were washed once with ice-cold phosphate-buffered saline (PBS), and then lysed in 200 μl icecold lysis buffer (50 mM Tris-HCl pH 7.4, 150 mM NaCl, 1 mM EDTA, 1% v/v Triton X-100, 10%v/v glycerol, 1x cOmplete protease inhibitor cocktail (Roche), and 1x PhosSTOP (Roche)). Cells were scraped off, collected in an eppendorf tube, and rotated for 30 min at 4°C. Debris was pelleted at 12,000 g for 10 min at 4°C, and supernatant was removed to a new tube on ice. Protein concentration was normalized to 15 μg total protein per lane using Bio-Rad Protein Assay Dye. A 5x Laemmli loading buffer (250 mM Tris-HCl pH 6.8, 30% glycerol, 0.25% bromophenol blue, 10% SDS, 5% β mercaptoethanol) was added to each sample to 1x, and samples were denatured at 95°C for 5 min, then spun down. Wells of AnyKd Mini-PROTEAN TGX precast protein gels (Bio-Rad) were loaded with equal amounts of total protein, in between Precision Plus Dual Color protein ladder (BioRad). After electrophoresis, proteins were transferred onto a nitrocellulose membrane, and then blocked for 2 h at room temperature in PBS with 0.1% Tween (PBS-T) + 3% milk (blocking buffer) while rocking. Primary antibody staining was performed with gentle agitation at 4°C overnight using the conditions outlined in Table 2. After washing four times in appropriate blocking buffer, secondary antibody staining was performed for 1 h at room temperature using antirabbit HRP or anti-mouse HRP secondary antibodies (Promega,1:10,000) in blocking buffer. Membranes were washed 3x in blocking buffer and then 1x in PBS-T without milk. Membranes were incubated with SuperSignal West Dura or Femto (Thermo Fisher Scientific) for 5 minutes. Membranes were imaged on a LI-COR Odyssey gel imager for 0.5–10 min depending on band intensity.

## Supporting information

Visualization of eIF2B conformational and Switch-Helix transformation. Switch-helix side-chain residues are visualized in red and ISRIB pocket is visu

## Acknowledgements

We thank Morgane Boone, Michael, Schoof, and the Walter and Marqusee labs for helpful discussions throughout the course of this project; Johanna Lindner for preliminary optimization of eIF2B HDX-MS coverage; Calico for a generous gift of purified eIF2 heterotrimer; and Z Yu and D Bulkley of the UCSF Center for Advanced Cryo-EM facility, which is supported by NIH grants S10OD021741 and S10OD020054 and the Howard Hughes Medical Institute (HHMI). We also thank the QB3 shared cluster for computational support.

## Funding

This work was supported by generous support from Calico Life Sciences LLC (to PW), a generous gift from The George and Judy Marcus Family Foundation (To PW), the Jane Coffin Child Foundation Postdoctoral Fellowship (to RL), Helen Hay Whitney Postdoctoral fellowship and 1K99GM143527 (to SS) and the Damon Runyon Cancer Research Foundation Postdoctoral fellowship (to LW). SM is supported by the Chan Zuckerburg Bi-Hub and NIH grant GM050945. PW was an investigator with the Howard Hughes Medical Institute.

## Author contributions

Conception and design: R Lawrence, S Marqusee, P Walter. Acquisition of data: R Lawrence, S Shoemaker, A Deal, S Sangwan, A Anand, L Wang. Analysis and interpretation of data: R Lawrence, S Shoemaker, A Deal, L Wang, S Marqusee, P Walter. Writing (original draft): R Lawrence, A Deal. Writing (review and editing): R Lawrence, S Shoemaker, A Deal, S Sangwan, A Anand, L Wang, S Marqusee, P Walter.

## Conflict of interest statement

PW is an inventor of ISRIB, patent held by the Regents of the University of California that describes ISRIB and its analogs. Rights to the invention have been licensed by UCSF to Calico. For the remaining authors, no competing financial interests exist.

---

**Supp. Fig. 1.**
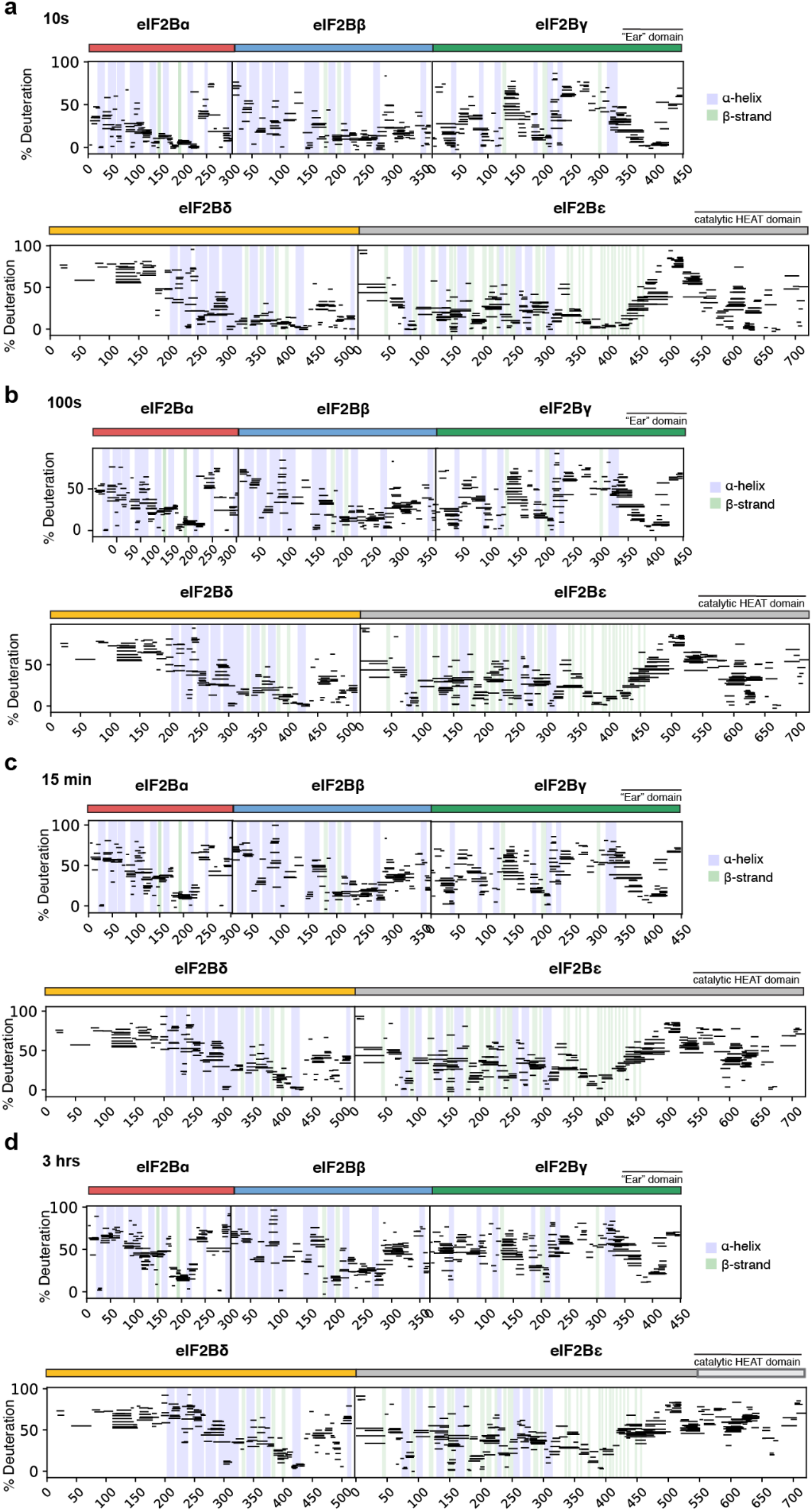
eIF2B deuteration over time. Percent deuteration after (**a**) 10 seconds (**b**) 100 seconds (**c**) 15 minutes (**d**) 3 hours of deuterium exchange for every peptide in one apo eIF2B dataset. Solid colored bars indicate each eIF2B subunit, corresponding to color scheme in Fig. 1. Each horizontal line represents an individual peptide spanning the residues indicated on the x axis, with percent deuteration (neglecting back exchange) indicated on the y axis. α-helices are indicated in blue vertical lines, and β-strands are indicated in green vertical lines, derived from apo eIF2B structure PDB 7L70.

**Supp. Fig. 2.**
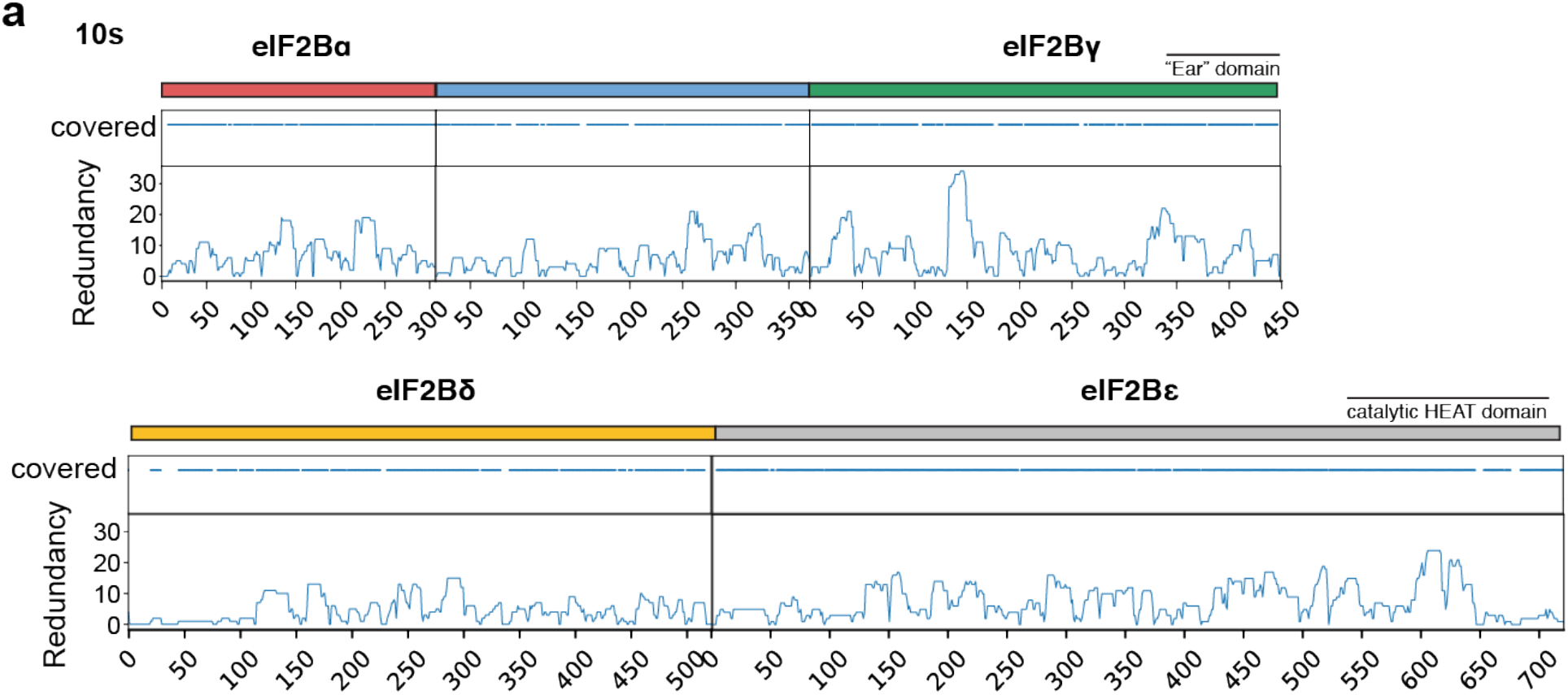
Coverage and redundancy results from HDX-MS experiments. (**a**) coverage and redundancy at each eIF2B residue for peptides included in all conditions of the comparative apo eIF2B vs NSs-bound eIF2B vs eIF2-P-bound eIF2B experiment. Solid colored bars indicate each eIF2B subunit, corresponding to color scheme in Fig. 1.

**Supp. Fig. 3.**
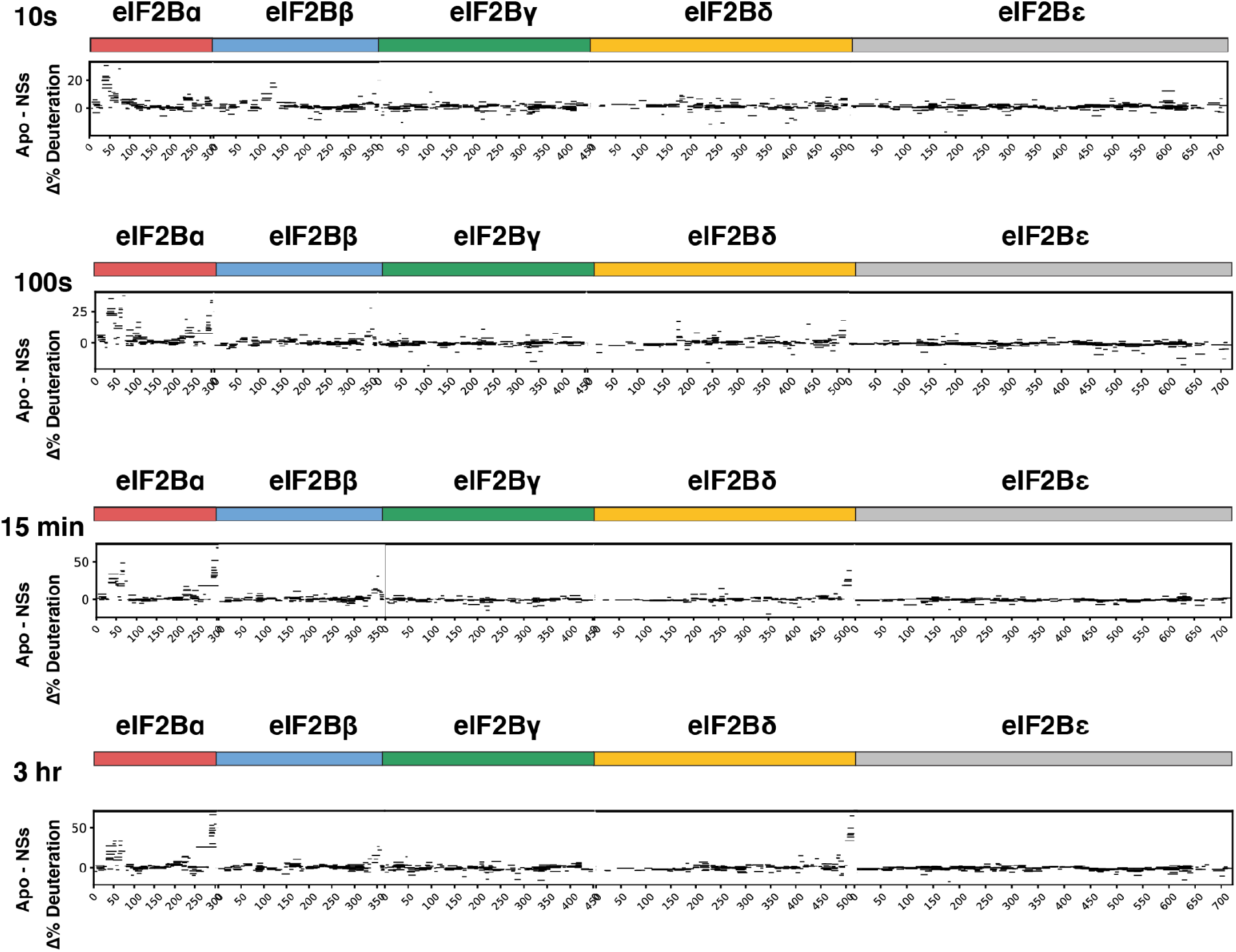
Representative percent deuteration difference maps over time for apo versus NSs-bound eIF2B(αβγδε)_2_ decamers. Positive values represent peptides that took up less deuterons in the NSs-bound state. Solid colored bars indicate each eIF2B subunit, corresponding to color scheme in Fig. 1.

**Supp. Fig. 4.**
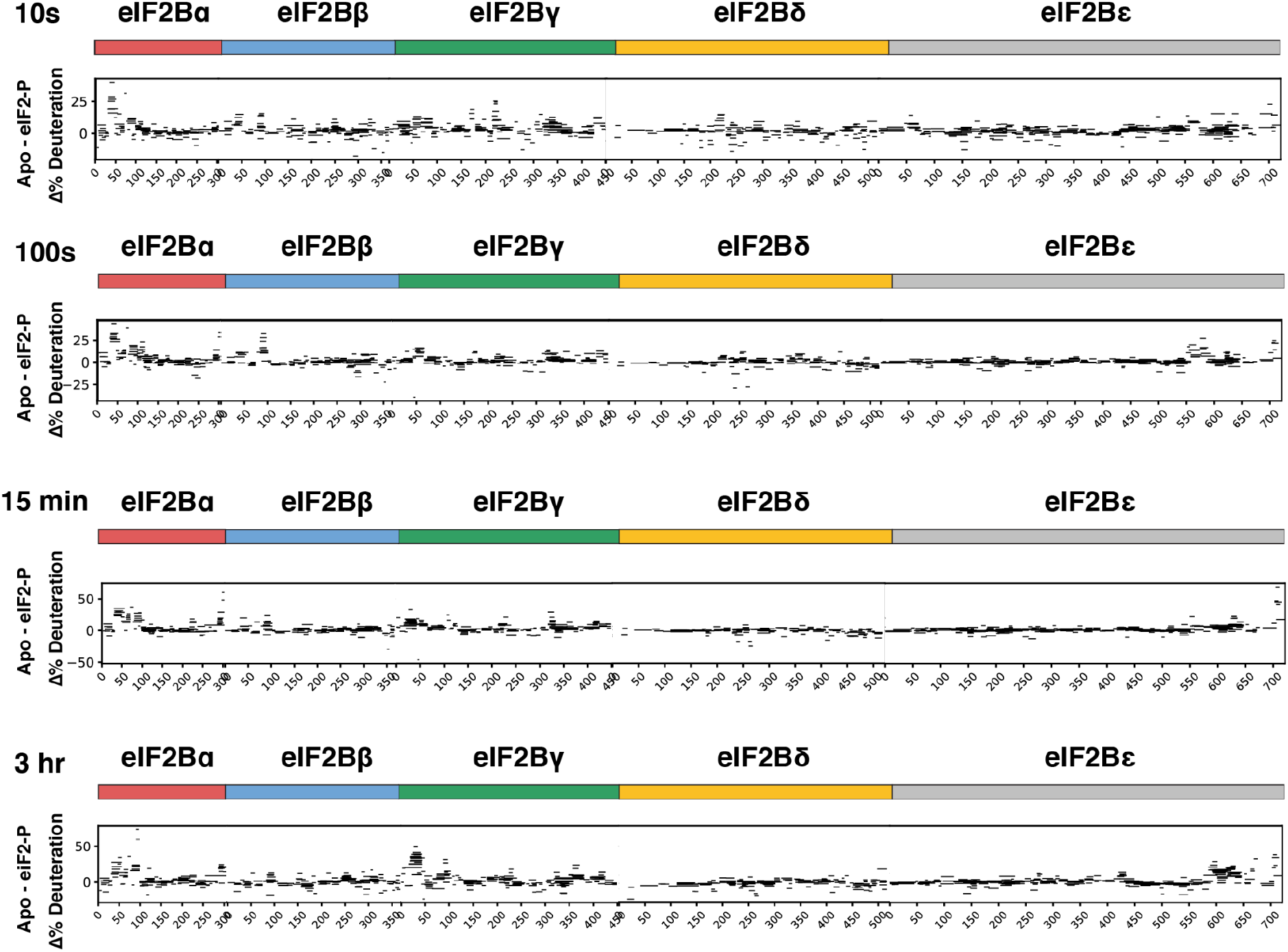
Representative percent deuteration difference maps over time for apo versus eIF2-P-bound eIF2B(αβγδε)2 decamers. Positive values represent peptides that took up less deuterons in the eIF2-P-bound state. Solid colored bars indicate each eIF2B subunit, corresponding to color scheme in Fig. 1.

**Supp. Fig. 5.**
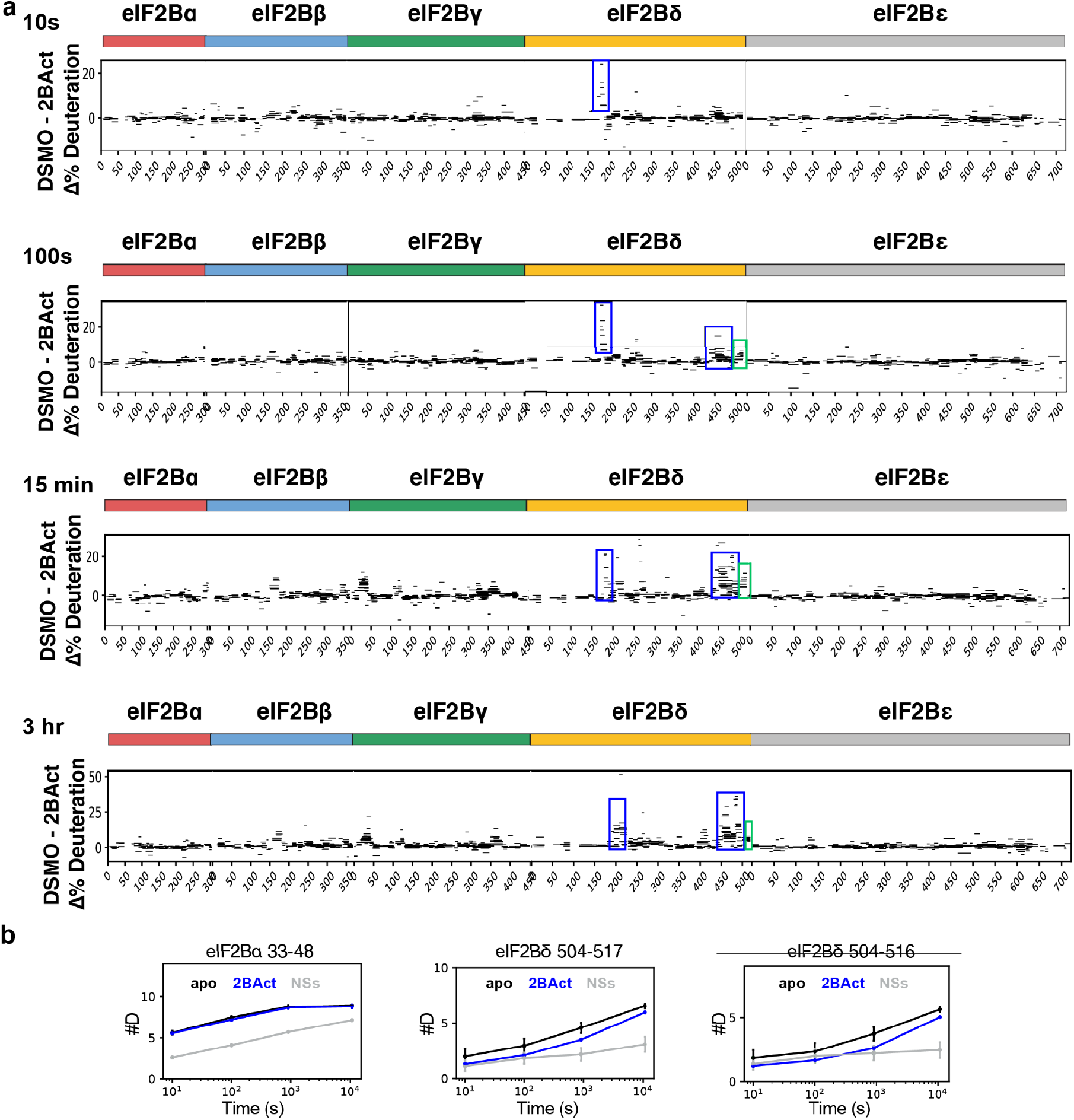
HDX-MS measurement of eIF2B protection by 2BAct. (**a**) Representative percent deuteration difference maps at the indicated timepoints for apo versus 2BAct-bound eIF2B(αβγδε)_2_ decamers. Positive values represent peptides that took up less deuterons in the 2BAct—bound state. Solid colored bars indicate each eIF2B subunit, corresponding to color scheme in Fig. 1. (**b**) Representative peptide uptake plots comparing 2BAct-bound (blue) vs NSs-bound (grey) vs apo eIF2B (black). The total number of exchanged deuterons per condition are plotted over time (representing an average of three independent experiments) ± SD.

**Supp. Fig. 6.**
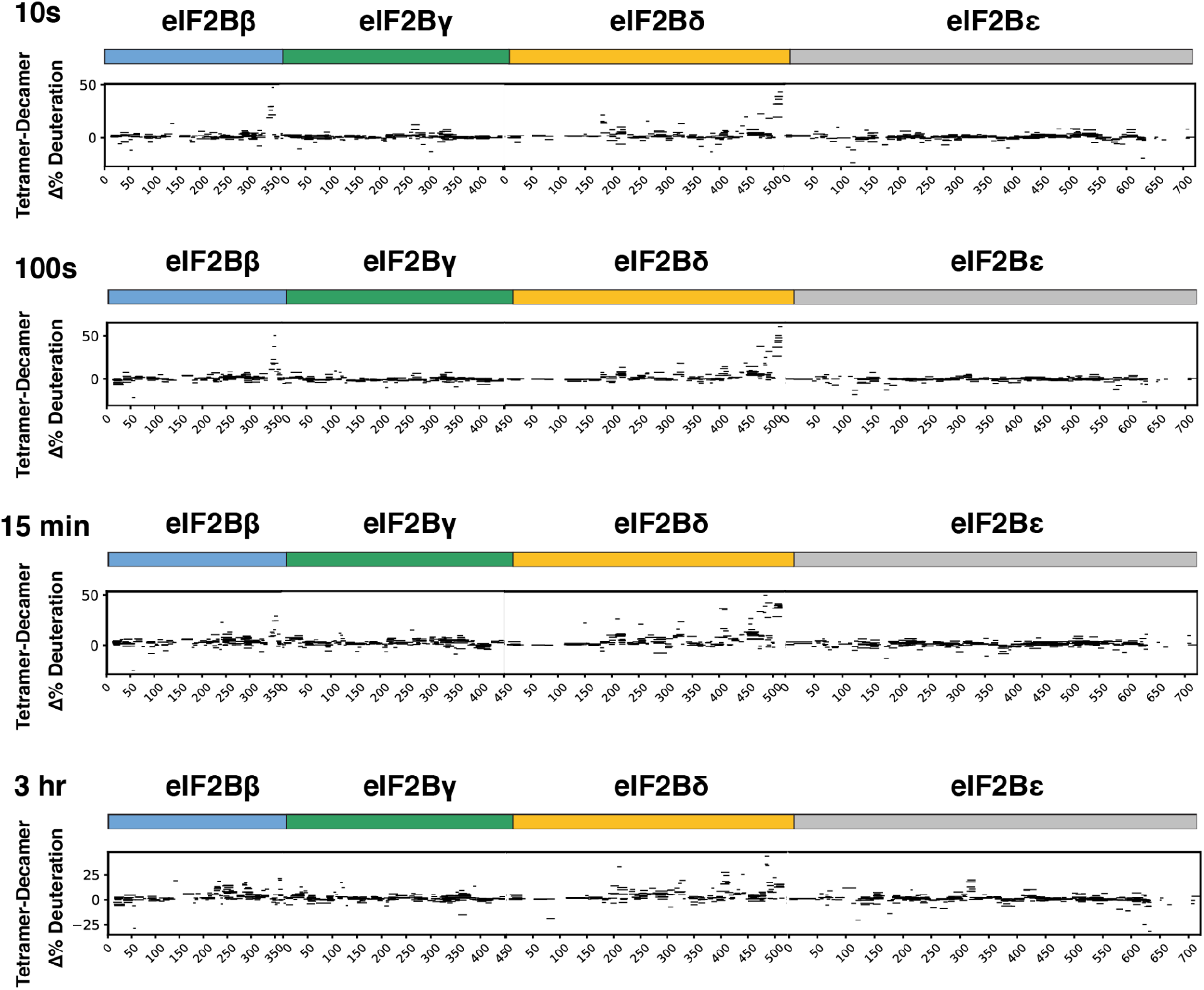
Representative percent deuteration difference maps at the indicated timepoints for eIF2Bβγδε tetramer versus eIF2B(αβγδε)_2_ decamer. Solid colored bars indicate each eIF2B subunit, corresponding to color scheme in Fig. 1. Positive values represent peptides that took up less deuterons in the decameric state.

**Supp. Fig. 7.**
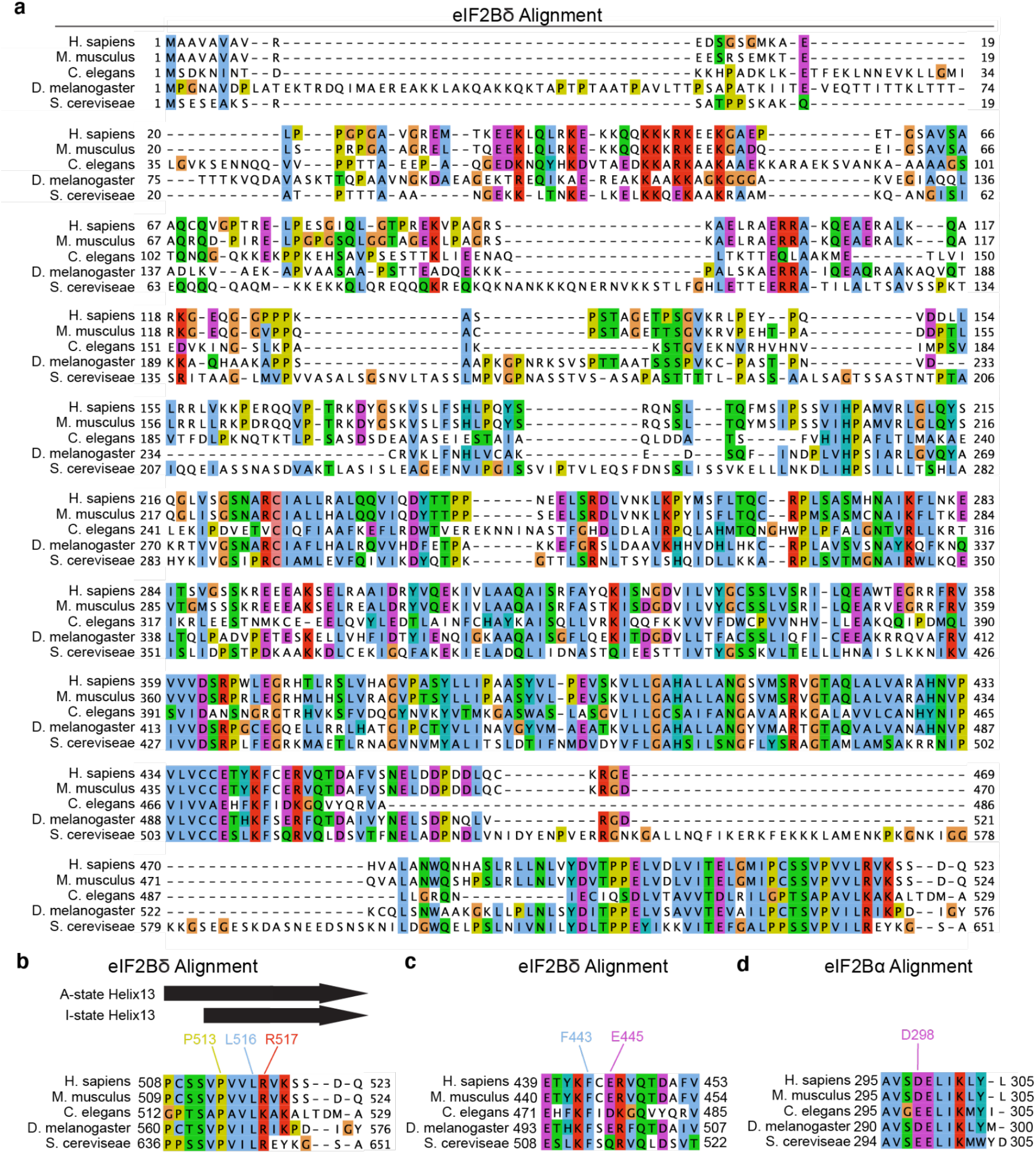
Sequence conservation of eIF2B Switch-Helix residues. (**a**) Multiple sequence alignment of eIF2Bδ from indicated organisms, with Clustal color-coding. (**b**) Multiple sequence alignment of eIF2B Switch-Helix from indicated organisms. Secondary structure is indicated above. (**c**) Multiple sequence alignment of eIF2Bδ residues that interact with the eIF2B Switch-Helix from indicated organisms. (**d**) Multiple sequence alignment of eIF2Bα residues that interact with the eIF2Bδ Switch-Helix from indicated organisms.

**Supp. Fig. 8.**
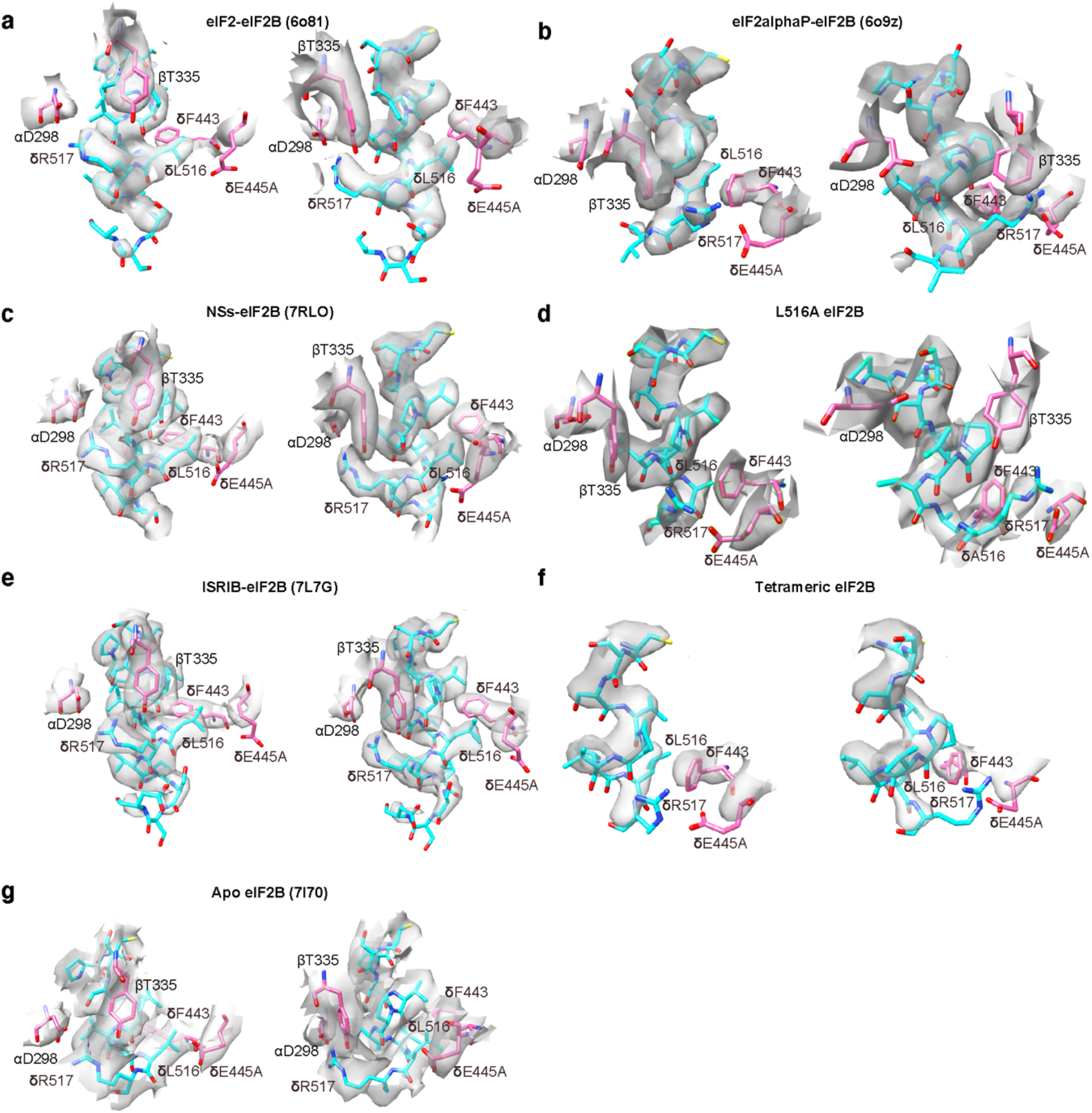
EM density surrounding eIF2Bδ Switch-Helix side-chains. Shown are EM density and atomic model of two representative views of relevant side-chains in the following eIF2B states: (a) eIF2-bound decameric A-state (PDB 6o81, EMD0649), (b) the eIF2αP-bound decameric I-state (PDB 6o9z, EMD0664), (c) the NSs-bound decameric A-State (PDB 7RLO, EMD24235), (d) the L516A decameric I-State (PDB TBD, EMD TBD), (e) the ISRIB-bound decameric A-State (PDB 7L7G, EMD7443), (f) the tetrameric state (PDB TBD, EMD TBD), (g) the decameric Apo eIF2B A-like state (PDB 7L70, EMD23209). Density zones were set to a radius of 2.0Å except for (a) which was 2.5Å and (f) which was 2.4Å.

**Supp Fig. 9.**
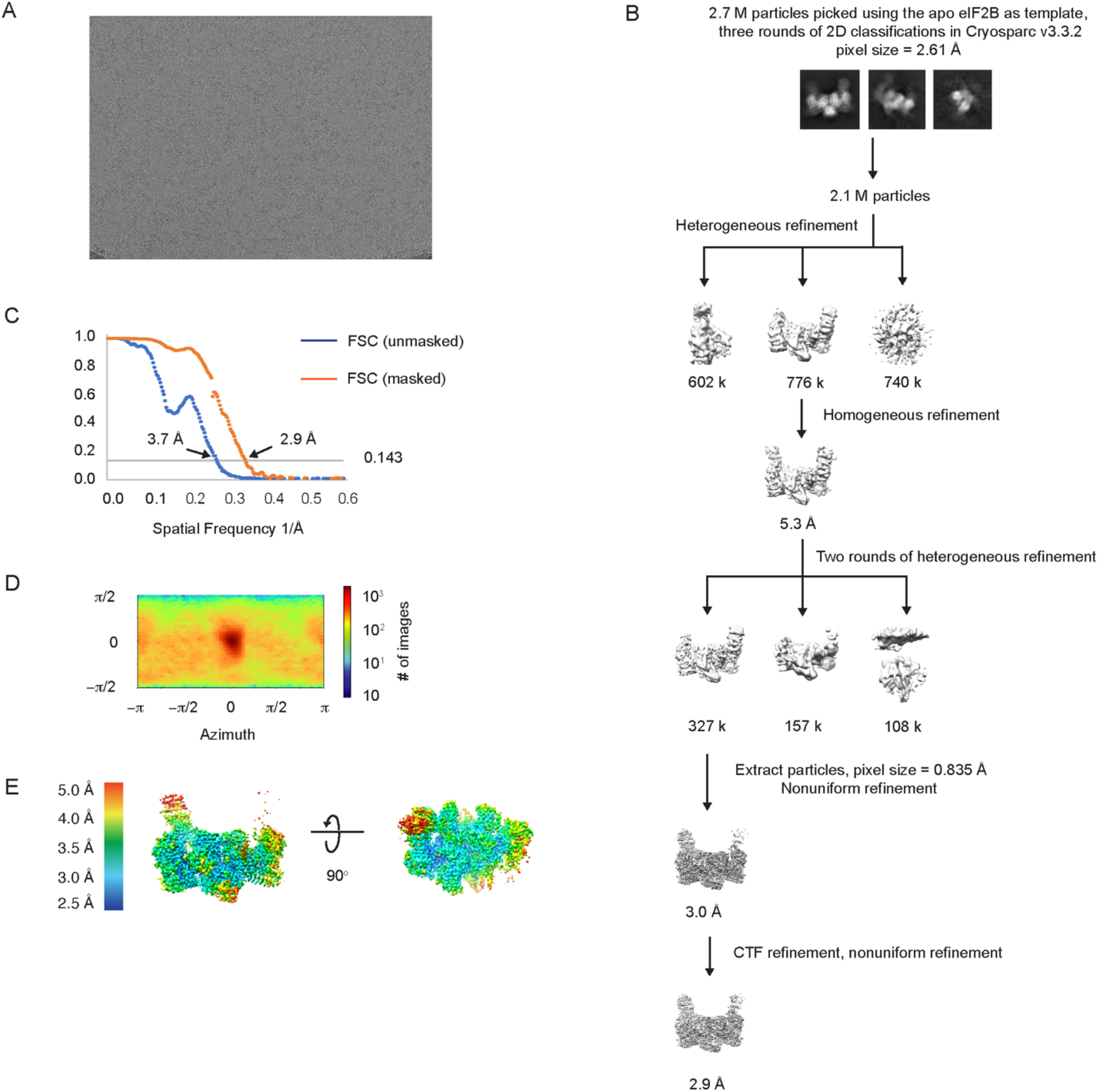
Cryo-EM data analysis of the eIF2B^δL516A^ structure. (**a**) Representative micrograph showing the quality of data used for the final reconstruction of the eIF2B^δL516A^ structure. (**b**) Data processing scheme of the eIF2B^δL516A^ structure. (**c**) Fourier shell correlation (FSC) plots of the 3D reconstructions of eIF2B^δL516A^ unmasked (dark blue), masked (orange). (**d**) Orientation angle distribution of the eIF2B^δL516A^ reconstruction. (**e**) Local resolution map of the eIF2B^δL516A^ structure.

**Supp. Fig. 10.**
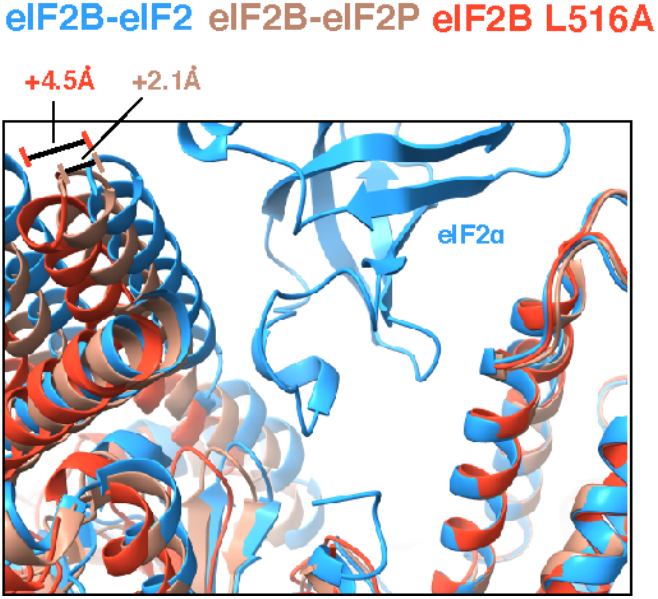
eIF2B L516A mutation widens eIF2α binding pocket more than eIF2αP binding. Overlay of atomic models of eIF2B-eIF2 (blue, PDB 6o81), eIF2B-eIF2αP (light peach, PDB 6o9z), and eIF2B L516A (orange) showing widening of eIF2α binding pocket by 2.1 Å for eIF2B-eIF2αP and 4.5 Å for eIF2B L516A.

**Supp. Fig. 11.**
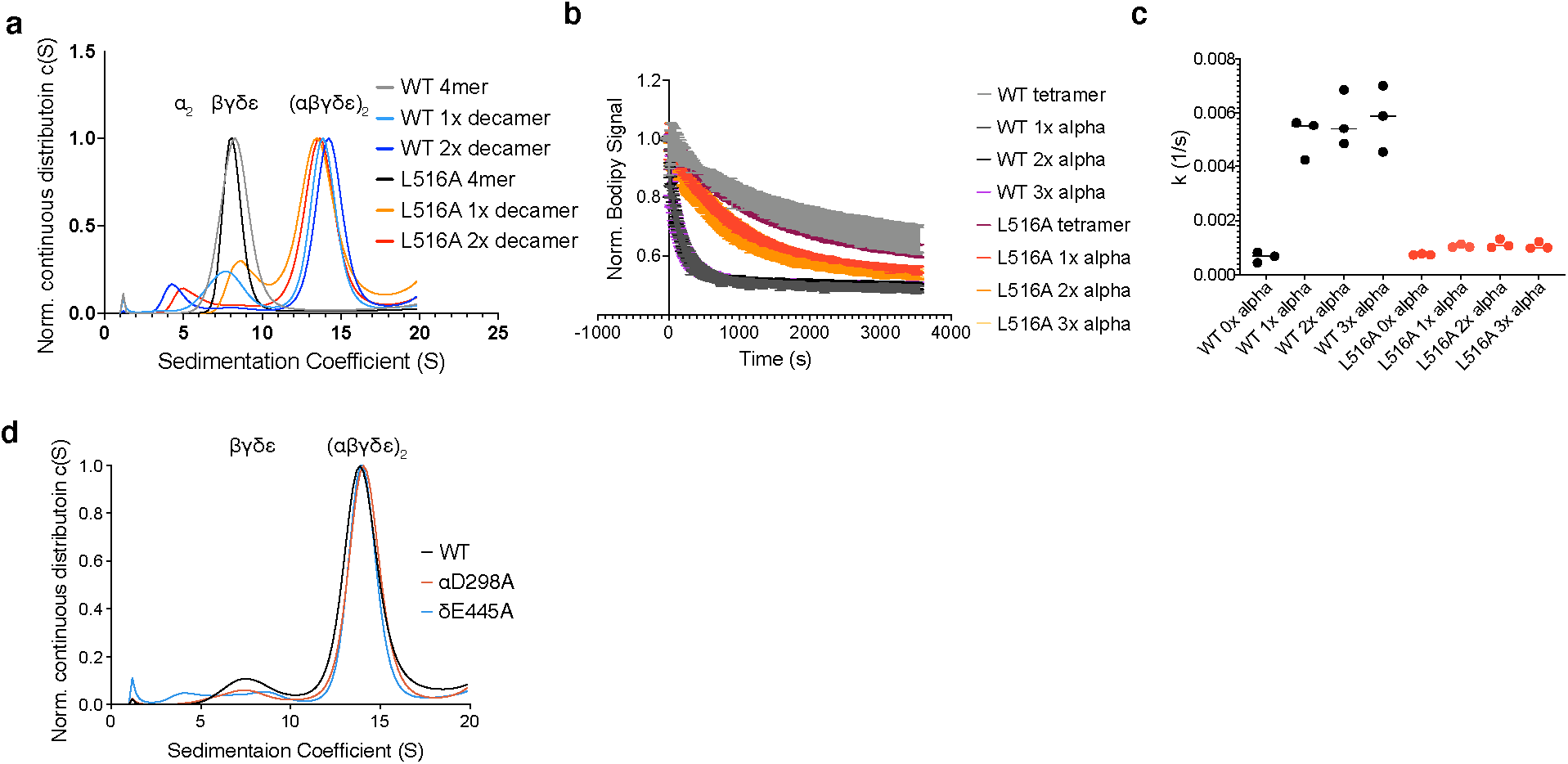
Decamerization propensity of eIF2B Switch-Helix variants. (**a-c**) eIF2B δL516A variant has a slight decrease in decamerization propensity that can be overcome by superstoichimetric eIF2Bα_2_. For all, “1X decamer” was assembled with a 2:1 stoichiometry of eIF2Bβγδε tetramer:eIF2Bα_2_ dimer. “2x decamer” was assembled with a 1:1 stoichiometry of eIF2B tetramer:eIF2Bα_2_ dimer. “3x decamer” was assembled with a 1:1.5 stoichiometry of eIF2Bβγδε tetramer:eIF2Bα_2_ dimer. (**a**) Sedimentation velocity analytical ultracentrifugation analysis of eIF2B(αβγδε)_2_ WT and δL516A decamer assembled with varying stoichiometries of eIF2Bα_2_ dimer. (**b**) Bodipy-GDP nucleotide loading assay of eIF2B(αβγδε)_2_ decamers (final concentration 5 nM) with the indicated point mutation and eIF2Bα_2_ stoichiometry. Shown are averages and S.E.M. for three experimental replicates. (**c**) Calculated rate constants for Bodipy-GDP nucleotide loading assay, as in (b). (**d**) Sedimentation velocity analytical ultracentrifugation analysis of eIF2B(αβγδε)_2_ decamers with indicated point mutations.

**Supp. Fig. 12:**
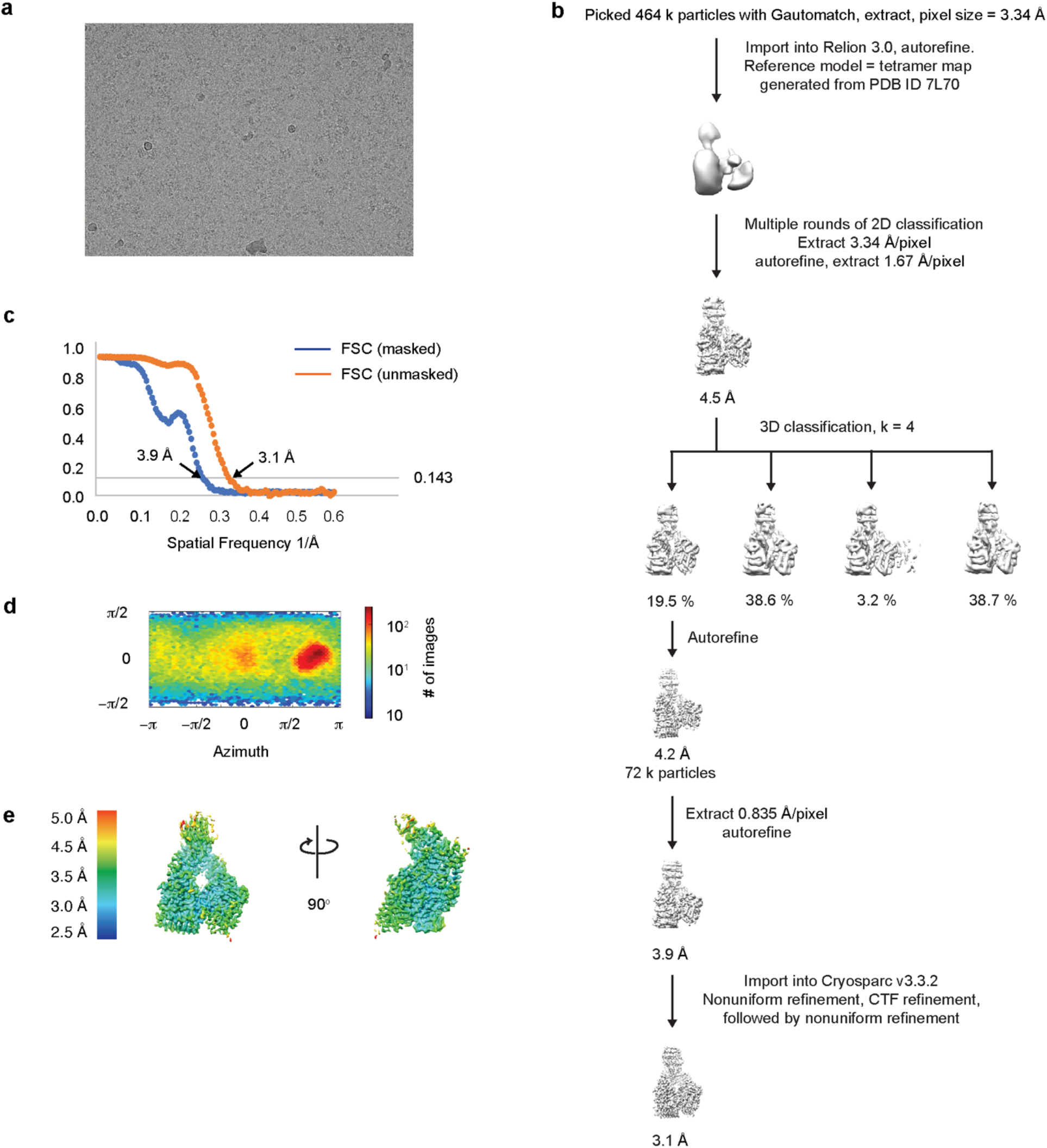
Cryo-EM data analysis of the tetrameric eIF2Bβγδε structure. (**a**) Representative micrograph showing the quality of data used for the final reconstruction of the eIF2Bβγδε structure. (**b**) Data processing scheme of the eIF2Bβγδε structure. (**c**) Fourier shell correlation (FSC) plots of the 3D reconstructions of eIF2Bβγδε unmasked (dark blue), masked (orange). (**d**) Orientation angle distribution of the eIF2Bβγδε reconstruction. (**e**) Local resolution map of the eIF2Bβγδε structure.

**Supp. Movie 1: Visualization of eIF2B conformational and Switch-Helix transformation.**

Switch-helix side-chain residues are visualized in red and ISRIB pocket is visualized in green.

